# Scalable joint non-negative matrix factorisation for paired single cell gene expression and chromatin accessibility data

**DOI:** 10.1101/2023.09.25.559293

**Authors:** William Morgans, Andrew D. Sharrocks, Mudassar Iqbal

**Affiliations:** Division of Molecular & Cellular Function, Faculty of Biology, Medicine and Health, University of Manchester, Manchester M13 9PL, UK; Division of Informatics, Imaging and Data Sciences, Faculty of Biology, Medicine and Health, University of Manchester, Manchester M13 9PL, UK

**Keywords:** Single cell multi-omics, Joint non-negative matrix factorisation, Topic modelling, Transcriptional regulation

## Abstract

Single cell multi-modal technologies provide powerful means to simultaneously profile components of the gene regulatory pathways of individual cells. These are now being employed to study gene regulatory mechanisms in a variety of biological systems. Tailored computational methods for integration and analysis of these data are much-needed with desirable properties in terms of efficiency -to cope with high dimensionality of the data, inter-pretability -for downstream biological discovery and hypothesis generation, and flexibility -to be able to easily incorporate future modalities. Existing methods cover some but not all of the desirable properties for effective integration of these data.

Here we present a highly efficient method, intNMF, for representation and integration of single cell multi-modal data using joint non-negative matrix factorisation which can facilitate discovery of linked regulatory topics in each modality. We provide thorough benchmarking using large publicly available datasets against five popular existing methods. intNMF performs comparably against the current state-of-the-art, and provides advantages in terms of computational efficiency and interpretability of discovered regulatory topics in the original feature space. We illustrate this enhanced interpretability in providing insights into cell state changes associated with Alzheimer’s disease. intNMF is available as a Python package with extensive documentation and use-cases at https://github.com/wmorgans/quick_intNMF

## Introduction

High throughput single cell ‘omics technologies have emerged as important tools in studying single cell states in complex tissues, ranging from developmental studies to complex diseases such as cancer. Recently technologies have been developed that enable simultaneous omics measurements in single cells [1]. For example, methylome and transcriptome [2]; chromatin and transcriptome [3, 4]; as well as triplets, chromatin, transcriptome and surface proteins [5]. These technologies help reveal regulatory changes which drive different gene expression programs and hence better determine cellular identity. With 10X genomics releasing the first commercially available solution (Chromium Single Cell Multiome ATAC + Gene Expression) it is likely the amount of paired ATAC-RNA (chromatin-transcriptome) data will increase [6]. One recent paper identified factors determining lineage plasticity in prostrate cancer [7] and another identified genes and genomic regions associated with Alzheimer’s disease [8], both using 10X Multiome data. Given the broad applicability of these technologies, their uptake for studying a variety of biological systems is likely to increase significantly. Hence, new computational methods are required which jointly analyse single cell RNA and ATAC count matrices and help discover underlying gene regulatory networks. A key step in multi-modal single cell integrative analysis is finding a low-dimensional and informative joint embedding from multiple distinct sets of features which capture biologically interesting sources of variation [9]. This is important as it enables all modalities to be used in cell type identification. However, due to high noise levels and differences in features of different modalities this is a challenging task. Most existing multi-omic integration methods focus on bulk data or single cell experiments carried out on different cells from the same population [10]. There are a handful of methods for integration of parallel ‘omics measurements from single cells. These are either based on metric learning (machine learning to identify a distance metric between data points) to generate multi-modal cell similarity metrics or latent variable modelling/matrix factorisation (MF)(finding lower dimensional matrices whose product recreates the original matrix). Metric learning approaches include Schema [11] and weighted nearest neighbour (Seurat WNN) [12]. Schema applies a quadratic programming formulation to linearly weight a secondary modalities features to maximise agreement with the primary modality. WNN is part of the popular R single cell analysis package Seurat [12], it calculates cell specific modality weighting which is used to create a joint nearest neighbour graph. The cell specific modality weighting is based on a cell’s similarity to its neighbours in a given modality. These methods are fast but the joint embedding is a cell-cell similarity matrix and therefore large and sparse. Importantly, these methods do not link measured features to the embedding and are therefore less interpretable. MF based approaches include multi-omic factor analysis (MOFA+) [13, 14], scAI [15], scREG [16], LIGER [17, 18], MultiVI [19] and MOJITOO [20]. MOFA+ performs probabilistic MF using Bayesian group factor analysis with variational inference scheme for the updates. This method produces interpretable results however takes much longer to run than gradient de-scent based methods. MultiVI also performs probabilistic MF using a variational autoencoder. This facilitates the use of a more complex model. However as the method is nonlinear, there is no direct link between the features and the joint low dimensional embedding. This method is primarily focused on removing batch effects and not understanding biology. MOJITOO uses Canonical Correlation Analysis (CCA) to create a joint embedding by linearly weighting separate low dimensional embedding to maximise their correlation. This method is computationally efficient but the factors of the joint embedding are constructed from modality specific embedding not the original features making them less interpretable.

Non-negative Matrix Factorisation (NMF) can perform fast and interpretable dimensionality reduction, enabling the speed of metric learning approaches to be combined with the interpretability of MF. NMF based methods have been successfully applied to bulk multi-omics analysis [21, 22], uni-modal single cell analysis [23] and unpaired single cell multi-omics (*i*.*e*. not measured in the same cells) analysis [17, 24, 25] and very recently paired multiomic analysis [15, 16]. scAI [15] applies NMF with a smoothed epigenetics matrix (*i*.*e*. pseudo-bulking ATAC data for inference). LIGER requires calculation of gene scores from chromatin accessibility data removing intergenic regions which are informative and often highly cell-type specific. scREG is also NMF based, however it introduces a third matrix based on genomic region and gene pairs (called *cis*-regulatory potential matrix). These methods rely on multiplicative updates to perform gradient descent for factorisation. Other update schemes exist which converge faster and more reliably [26]. More recently a method called *Mowgli* has been developed, which uses optimal transport to optimise a joint NMF model, however the authors do not recommend running the method on datasets with more than 1000 cells without GPUs [27]. Additionally both scAI and scREG require the entire dataset to be loaded into memory limiting the size of dataset which they can be applied to.

Here we propose an NMF based method, called intNMF, for the dimensionality reduction and exploratory analysis of paired multiomic single cell data. It implements a novel joint factorisation scheme, with no transformations leading to loss in chromatin accessibility information. It is computationally efficient in terms of memory and time allowing it to scale to large numbers of cells and features. intNMF generates an interpretable joint embedding to represent cells from multiple modalities which can be useful in downstream analysis. The method is implemented in Python and is compatible with the Python single cell ecosystem (*e*.*g*. Scanpy/anndata/MUON [28–30]). Given this, it is also easy to use R for certain downstream analysis due to ease of conversion between anndata objects and Seurat objects with the “SeuratDisk” R package or “anndata2ri” python package.

We quantitatively evaluate intNMF against Seurat WNN, scAI, scReg, MOFA+ and MOJITOO on two multiome datasets containing paired gene expression and chromatin accessibility data. intNMF and MOFA+ are then applied to a third dataset to compare the biological interpretability of the learned low dimensional embeddings and perform down-stream biological analysis of the topics.

## Methods

### intNMF

NMF is a linear dimensionality reduction method which approximates a non-negative matrix as the product of two lower dimensional (or low rank) non-negative matrices which minimise the reconstruction error, *i*.*e*.

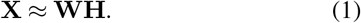

In the case of a scRNA-seq dataset: 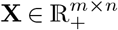 is a nonnegative cell by gene matrix to be factorised (*m* is the number of cells and *n* is the number of genes), a row of 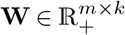 contains the coordinates of a single cell in terms of the nonnegative basis vectors given by 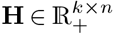. *K* is the desired number of dimensions for the factorisation, referred to here as topics due to NMFs similarity with topic modelling. A row in **W** describes a cell as a series of topic activities and which are quantitatively defined by gene scores (*i*.*e*. rows in **H**). *k* is selected such that *k << m* or *n*. The nonnegativity constraint has been found to produce a parts based representation of the original data [31]. The topic definitions (rows of **H**) contain real positive numbers like the original data, e.g., when NMF is applied to a collection of images each basis vector (or topic) is an image. These two qualities lead to interpretable and informative low dimensional embeddings. In the case of multiome (paired single cell RNA and ATAC) data, intNMF approximates two non-negative matrices so that they share a joint embedding (**W**) but have two sets of basis vectors (**H**), *i*.*e*.

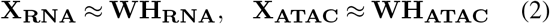

**W, H**_**RNA**_ and **H**_**ATAC**_ can be found by minimising a cost function describing the reconstruction error, *i*.*e*.

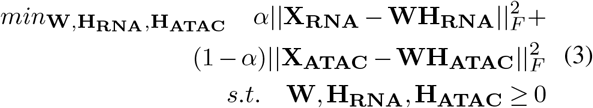

Here, the cost function used is the Frobenius norm, *α* allows the different modalities to be weighted differently in the cost function, by default it is set to 0.5. An element-wise l1 penalisation on **W** and **H** can be applied giving:

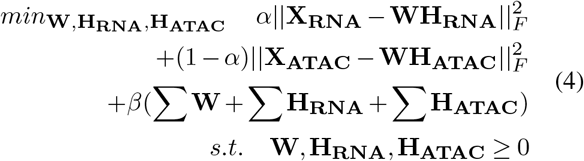

intNMF implements the accelerated hierarchical alternating least squares (acc-HALS) method [26], which we modified to jointly factorise two matrices. HALS is a block coordinate descent method where the optimisation problem is broken up into smaller sub-problems, in this case the columns of **W** (**w**_**k**_) and rows of **H** (**h**_**k**_). This is done by minimising the reconstruction error w.r.t a single topic on the residual between **X** and all the other topics. For example for the reconstruction of the RNA matrix:

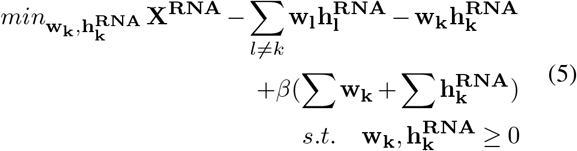

This results in the following update equations:

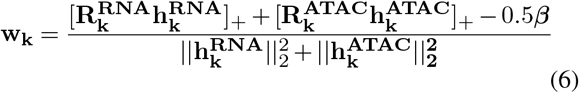

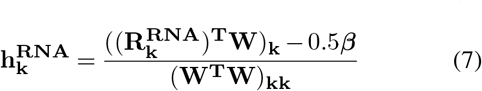

Note that **w**_**k**_ also depends on the ATAC modality which is omitted from equation 5 for clarity. 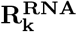 is the residual from reconstruction using the other columns of **W** and rows of **H** *e*.*g*.

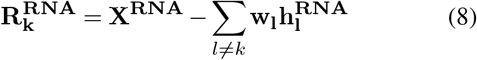

The equations for 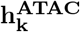 and 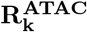 are identical to equa-tions 7 and 8 with the modality switched.

Subbing **R**_**k**_ back into these updates allows parts of the updates to be precomputed and reused for every vector. This can be seen in Algorithm 1. This gives the following update equations which are used by intNMF (Supplementary Algorithms 4 and 5)

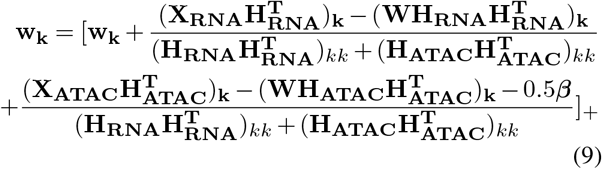

and

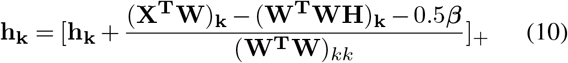

If the updates result in negative numbers they are replaced with small positive numbers to ensure the positivity constraint.

The acc-HALS algorithm updates **W** and **H** multiple times at each epoch, to take advantage of precomputing values which do not change. After updating all columns/rows of **W** or **H** the matrix is updated again if both of the following criteria are met:

1. epoch run time *<* (precompute time (*e*.*g*. **X**^**T**^**W** and **H**^**T**^**H**) + last iteration duration)*/*2
2. sum of squares of update to **W** or **H** at last iteration *>* 0.01*sum of squares of update to **W** or **H** at first iteration

Updates to a single matrix are stopped when either of the conditions are no longer met. In our experience matrices are updated roughly 5 times per epoch. A complete derivation of the intNMF updates can be found in the Supplementary Methods.

#### Algorithm 1

intNMF Overview. The implementation details of *W* _*HALS* and *H*_*HALS* are given in Supplementary Algorithms 4 and 5, respectively.

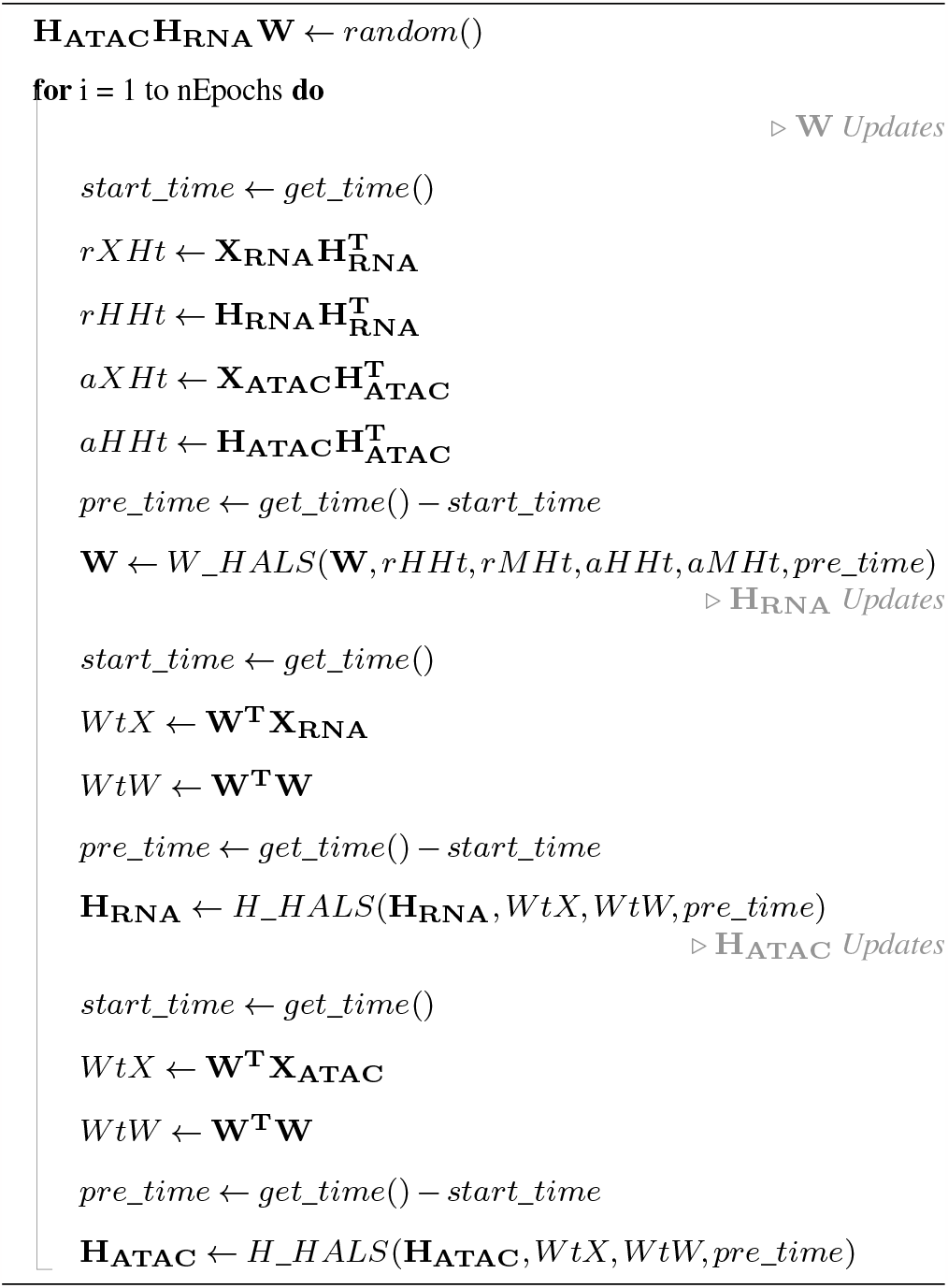

### Datasets

Two datasets were used to benchmark intNMF against existing methods and third as a narrative example of itnNMF’s capabilities. All datsets were generated with the 10x multiome kit and an overview of them is shown in Table 1. The first dataset (PBMC-Multiome) was generated from Human peripheral blood mononuclear cells (PBMC) taken from a female donor, and was published by 10x Genomics. The second dataset [32] (BMMC-Multiome) was generated from bone marrow mononuclear cells (BMMC) and includes data from 12 healthy human donors processed across multiple sites. This dataset was created for the Neurips 2021 Data Integration challenge and was purposefully designed to create batch effects [9]. A third dataset was taken from a recent publication of a post mortem study of samples taken from the dorsolateral prefrontal cortex (DLPFC) of Alzheimer’s positive (7 samples) and negative (8 samples) individuals [8].

**Table 1.** Characteristics of the datasets used for benchmarking and application of intNMF. Alzheimer’s data have 8 main cell types with 36 subtypes.

| Dataset | Method | Species | Organ | # cells | # cell types | # features (gene/region) |
| --- | --- | --- | --- | --- | --- | --- |
| PBMC-Multiome | Multiome | Human | PBMC | 11909 | 13 | 36601/108377 |
| BMMC-Multiome | Multiome | Human | BMMC | 69249 | 22 | 13431/116490 |
| Alzheimers-Multiome | Multiome | Human | DLPFC | 105332 | 8/36 | 36601/150614 |

### Preprocessing

All datasets used for benchmarking were subjected to the same basic preprocessing steps. For the gene expression matrices, genes were filtered by minimum and maximum number cells expressed in (*≥*1% and*≤*90% of total cells) and cells by minimum and maximum number of expressed genes (*≥* 5% and *≤* 80% of total genes) and mitochondrial reads (*≤* 25% of total reads).

For the chromatin accessibility matrices, regions were filtered by minimum and maximum number of cells accessible in (*≥* 1% and *≤* 80% of total cells) and cells were filtered by the number of accessible regions (*≤* 80%) .

As preprocessing can be a crucial step in determining a methods efficacy, method specific preprocessing was carried out inline with recommendations from each methods documentation, see Supplementary Methods for details. For the RNA modality this typically involved library sized normalisation followed by *log*(*x* + 1) transformation and for the ATAC modality Term Frequency-Inverse Document Frequency (TF-IDF) transforming the data. TF-IDF transformation is implemented in the intNMF package as:

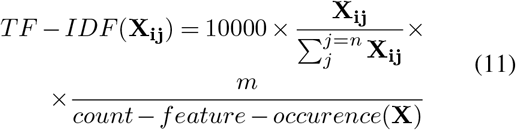

where **X** is the cell-feature count matrix (either chromatin accessibility or gene expression), *count−feature− occurence*(**X**) is the number of cells a region is accessible in, *n* is the number of features and *m* is the number of cells.For non-NMF methods matrices are often scaled to have zero mean and unit variance.

### Benchmarking

To compare the effectiveness of different methods, a range of metrics were used. The compactness of cell types in the joint embedding was calculated using the silhouette score. This measures the distance (measured in the joint embedding) between cells annotated with the same type relative to the other cell types.

Additionally the ability of the joint embedding to cluster cells was measured, either by generating cell-cell similarity matrices (from the joint embedding) which are then used to perform Leiden clustering or by a cell’s max topic/factor score. The clustering method which produces the best scoring metric (for a given dimensionality reduction method) is reported here. Cell clusters are compared to known labels and evaluated using normalised mutual information (NMI) [33] and Adjusted Rand Index (ARI) [34]. ARI measures the proportion of cells correctly clustered adjusted for chance. NMI, arises from information theory and describes how much information about the ground truth labels are obtained from the calculated clusters.

### Competing methods

A brief overview of the competing methods can be seen in Table 2, and a more thorough description of the methods, including details of their implementation, can be found in the Supplementary Methods.

**Table 2.** Competing methods summary.

| Name (version) | Language | Key Features |
| --- | --- | --- |
| MOFA+ [13] (1.2.2) | Python & R | MF - Probabilistic PCA via variational inference |
| MOJITOO [20] (1.0) | R | MF - CCA on PCA of RNA and LSI of ATAC |
| scAI [15] (1.0) | Python & R | NMF - multiplicative updates, smoothed ATAC by learnt nearest neighbour matrix |
| scREG [16] (0.1.0) | R | NMF - multiplicative updates, tri-modal factorisation, cis-regulatory potential matrix |
| Seurat [12] (4.2.0) | R | Graph based - multi-modal nearest neighbour analysis |

### Rank selection

The number of topics is an important hyperparameter to select. By default it is left to the user to select the value based on prior knowledge of the dataset. However intNMF also provides two functions for model selection. Both methods require multiple models with different numbers of topics to be fit to the data. The number of topics is then selected based on either Bayesian information criterion (BIC) or the knee point of the loss as function of topic number.

The first relies on a modified version of BIC for NMF [35], given below:

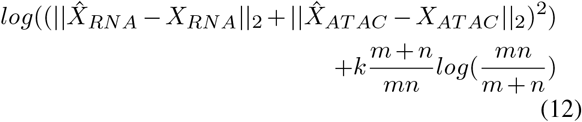

where 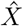 denotes the NMF reconstructed matrix of a given modality. Alternatively, intNMF uses kneedle [36] to identify where the reduction in loss flattens with the addition of further topics.

In practice the minimal value of BIC and the “knee” point in the function of loss vs topic number return similar values which in our experience roughly reflect the number of cells present in a system.

## Results

### Benchmarking computational methods for joint embedding of multiome data

We first benchmarked intNMF’s ability to generate an informative latent embedding against other published methods for joint analysis of single cell multi-omics data, using a publicly available 10X Multiome dataset of *∼*12, 000 PBMCs with ground truth cell types annotations. The quality of the embedding was measured by its ability to recapture the ground truth cell labels. This was done in two ways, firstly by performing clustering using the embedding and comparing this with the ground truth labels (MI and ARI metrics) and secondly by measuring how compact cells of the same type are in the joint embedding (silhouette score).

The benchmarking results on the PBMC-multiome data (12000 cells) are shown in Figure 1A. Two intNMF models are shown with l1-regularisation: intNMF-l1 (leiden) with k=35, epochs=75, and *β*=1 (following by Leiden clustering on W matrix) and intNMF-l1 (topic) with k=10, epochs=50, and *β*=20 (following max topic clustering on W matrix). We generally find that intNMF gives best clustering results with max topic with fewer topics and larger l1 regularisation and better clustering via Leiden clustering with more topics and lower l1 regularisation. One example of intNMF without l1-regularisation is given with k=15 and epochs=10. scAI was run without error message or warning however has been omitted from the figures as the returned cell embedding and feature loading matrices were filled with small positive numbers, potentially suggesting failed convergence. A UMAP generated from scAI’s embedding can be seen in Supplementary Figure 1.

**Fig. 1.**
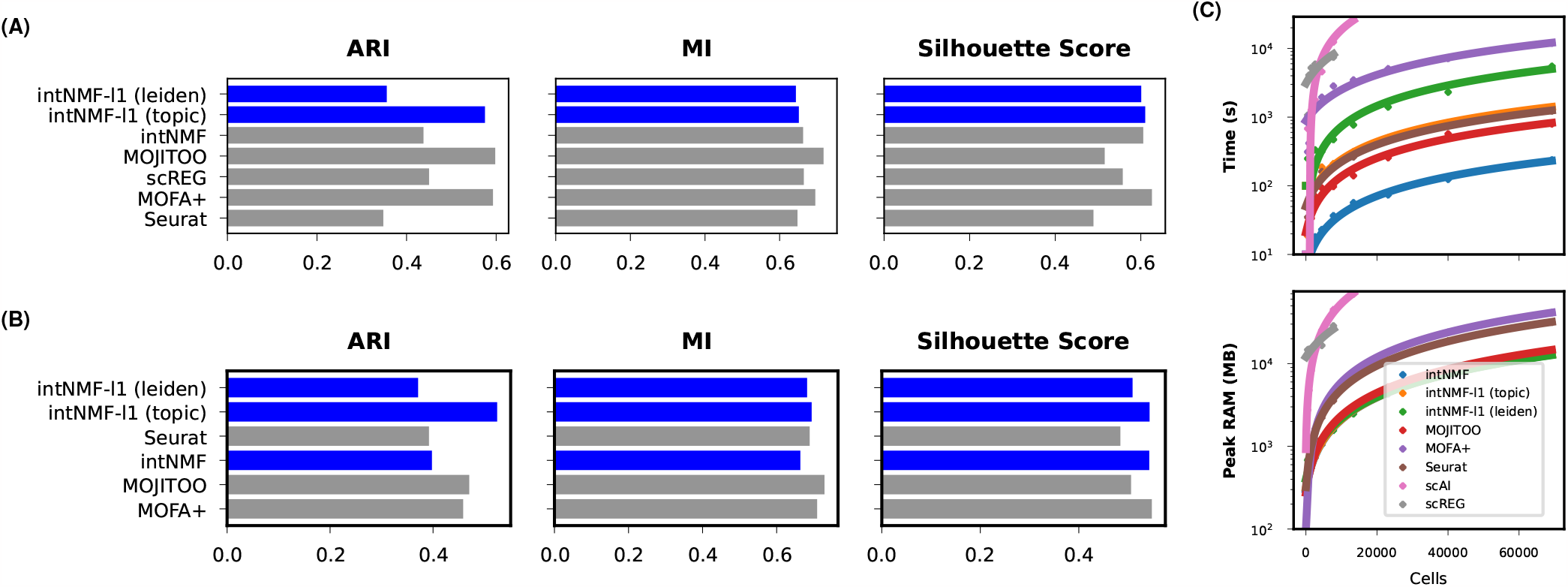
Benchmarking on PBMC Multiome. (A) Performance metrics based on clustering using different joint embeddings for the PBMC-Multiome dataset (*∼* 12, 000 cells). The performance metrics are Adjusted Rand Index (ARI, left), normalised Mutual Information (MI, centre) and Silhouette Score. (B) as in (A) but for the larger BMMC-Multiome dataset (69249 cells). (C) Process time and memory usage of the indicated methods on subsets of the BMMC-Multiome dataset.

MOJITOO, MOFA+ and intNMF-l1 (topic) have the best overall performance, when measured by ARI. All variants of intNMF perform comparably to other methods when measured via MI. However, intNMF (all variants) and MOFA+ both show superior performance, relative to the other methods, when measured by silhouette score (Figure 1A). UMAPs generated from each methods embedding can be seen in Supplementary Figure 1, with the ground truth cell types indicated by colour. The UMAPs share a similar structure and all the annotated cell types form distinct clusters in the UMAPS generated from the different methods, with the exception of scAI. Additionally similar cell types are typically found in adjacent clusters *e*.*g*. B Cell progenitor cells and pre B cells are closely juxtapositioned by the different methods. This indicates the methods can not only identify clusters and but also similarities between cell types in these clusters.

Next we benchmarked the methods on a larger dataset, the BMMC-Multiome, consisting of 69249 cells and 22 distinct cell types. scREG and scAI are omitted from this analysis as they encountered memory errors when run on this entire dataset. Two intNMF models are shown with l1-regularisation one with k=80, epochs=50, and *β*=0.1 (leiden) and k=22, epochs=75, and *β*=20 (topic). One example of intNMF without l1-regularisation is given where k=15 and epochs=10. intNMF-l1 (topic) has the best clustering performance measured by ARI, whereas MOJITOO performs best as measured by MI. intNMF-l1 (topic), intNMF and MOFA+ have comparable Silhouette Scores.

We compared the run time and memory requirements for all methods by sampling increasing number of cells from BMMC-Multiome data (Figure 1C). intNMF has similar run time and memory requirements to MOJITOO and Seurat, outperforming existing MF methods scAI, MOFA and scREG. This difference is most pronounced compared with the existing NMF methods scAI and scREG where results are only shown up to the point where they encounter memory errors (Figure 1C). intNMF’s run time and memory requirements are sensitive to the number of topics, k and epochs, increasing roughly linearly with both. Additionally, the run time of the methods depends on the number of features which they are applied to. MOJITOO acts on hundreds of features (separately low dimensional representations of the modalities), MOFA+ acts on thousands of selected highly variable features, whereas intNMF is applied to all features in the dataset (*i*.*e*. 13431 genes and 116490 regions). This has the downside of reducing the speed of the method but the benefit that topics are linked to all features in the original data.

Collectively, this benchmarking analysis shows that intNMF shows competitive performance in terms of clustering of cells (based on a combinations of metrics) and it is time and memory efficient compared to the state-of-the-art methods. A typical run of intNMF on 70000 cells takes approximately 5 minutes on a computer with 4 nodes with 16GB RAM each. Besides these performance measures, intNMF output is highly interpretable which greatly facilitates downstream analysis we show a detailed analysis through its application on brain data (multiple post-mortem samples, with/without Alzheimer’s disease) in the next section.

### Application to dorsolateral prefrontal cortex multiome data

To highlight intNMF’s ability to generate a biologically informative joint embedding and topic definitions, it was applied to a recently published 10x multiome dataset comprising *∼*100000 brain cells from the DLPFC from post mortem tissue of donors with and without Alzheimer’s [8]. We compared intNMF performance to the next best performing tool, MOFA+. For ease of comparison latent dimensions of both methods embedding are referred to as topics. intNMF was run with 25 topics as this was the number of topics which produced the lowest BIC value and was the elbow point of the loss as a function of number of topics. intNMF was also run with l1-regularisation with *β* = 20 as this level of regularisation performed well in the benchmarking and l1-regularisation should help the method learn informative features.

UMAPs generated from both intNMF and MOFA+ clearly resolve the annotated cell types in the dataset (Figure 2A). The topics generated by intNMF are more cell type-, or even sub cell type-specific, than the MOFA+ topics (Figures 2B and 2C). For example, intNMF topic 11 specifies astrocytes on the UMAP and topic 1 highlights a subset of the excitatory neurons (Figure 2B and Supplementary Figure 5D). In contrast, MOFA+ topics 4, 6 and 7 have positive and negative weights in the same cell type (Figure 2C and Supplementary Figure 4D). With the exception of topic 12, the intNMF topics are cell type-specific, whereas the MOFA+ topics have positive weights in multiple cell types (Figure 2D). intNMF topics are active in similar cell types which might be expected to share gene expression programs *e*.*g*. topic 5 for inhibitory and excitatory neurons. A more detailed analysis of the intNMF-derived topic contributions to each cell in the dataset can be seen in Figure 2E. It further illustrates the specificity of topic-mediated cell designations.

**Fig. 2.**
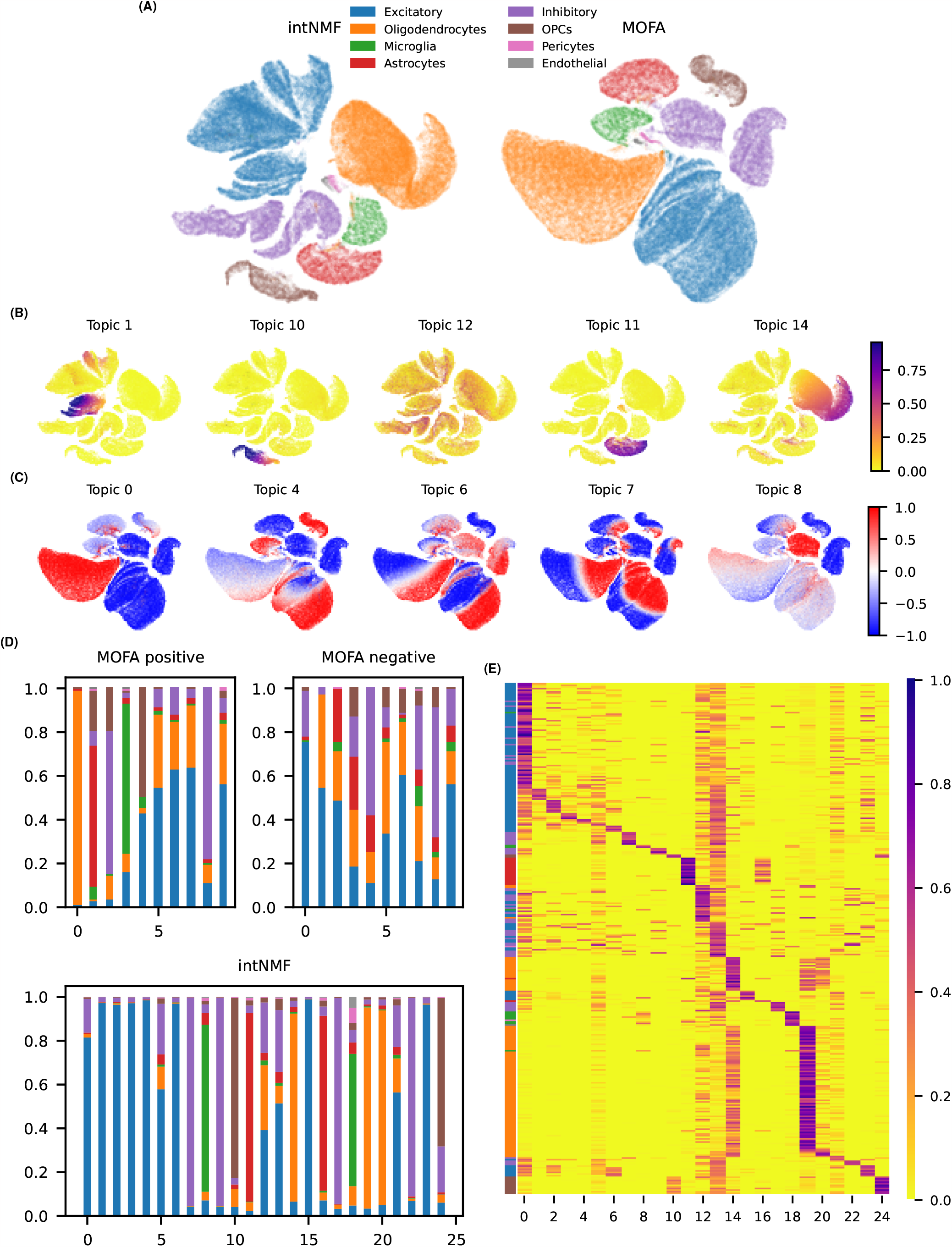
Comparison of intNMF and MOFA+ analysis of an Alzheimer’s disease dataset. (A) UMAP generated from nearest neighbour matrices using MOFA+ and intNMF embedding’s on the Alzheimer’s dataset annotated with cell type. (C) intNMF UMAP with a selection of topics scores shown (B). MOFA+ UMAP with a selection of topic scores shown. (D) Proportion of topic weights (*i*.*e*. values in **W**) contributing to the different cell types. The x-axis is topics and the y-axis shows the proportion of the sum of the topics weight is found is a given cell type. The cell type is indicated by colour following the same scheme as (A). (E) intNMF **W** matrix sorted by max topic with the cell type annotated on the leftmost column (colour scheme as in (A)).

While both MOFA+ and intNMF readily identify cell types and sub cell types (Supplementary Figures 11B and 10C), differences between Alzheimer’s positive and negative cells are harder to elucidate (Supplementary Figure 4A and 5A). This parallels the findings of the original study where Seurat WNN was applied for the joint analysis and did not resolve distinct Alzheimer’s clusters.

### intNMF topics are biologically relevant and comprise related RNA and chromatin features

Having established the ability of intNMF’s cell-topic matrix **W** to identify cells and sub-populations of cells, we next focused on investigating whether the learnt features which define topics are biologically relevant. The genes and regions which are important in defining topics are sparse, and this is reflected in the weights of intNMFs loading matrices *i*.*e*. **H** matrices. The majority of the weights are almost zero (Figure 3A). We also looked at relative contributions of individual modalities towards the joint topics, as it makes sense that certain biological topics should be driven more by RNA features, others by ATAC, or by a mixture of both. The topic loadings (*i*.*e*. rows of **H**_**RNA**_ and **H**_**ATAC**_) can be dominated by a single modality (*e*.*g*. topic 11 -RNA and topic 1 -ATAC) or equally contributed to by both (e.g., topic 13) (Figure 3A, Supplementary Figure 7).

**Fig. 3.**
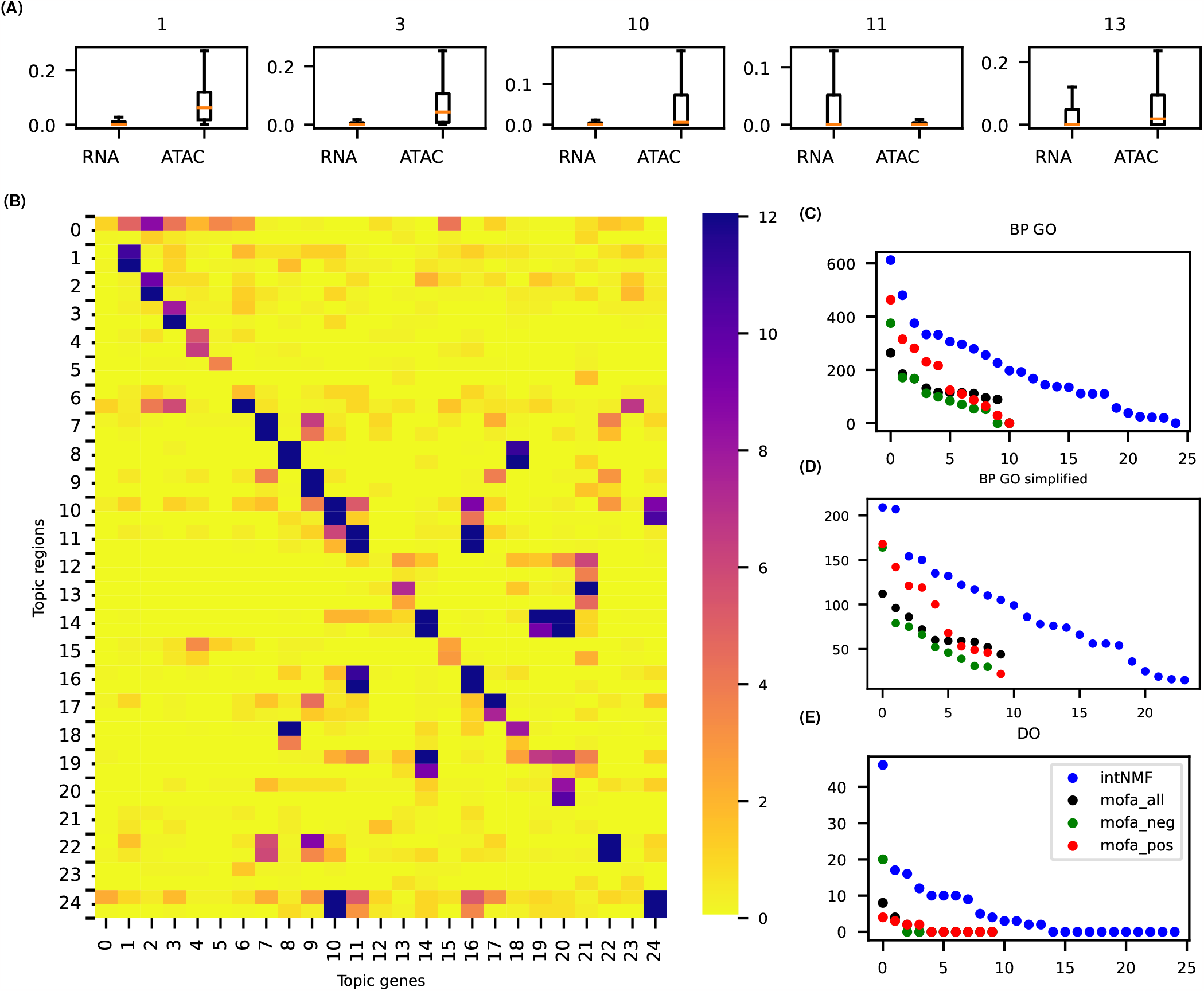
(A) Box plots of a subset of intNMF topic feature scores *i*.*e*. the values in rows of **H**_*AT AC*_ and **H**_*RNA*_. (B) PEGS analysis [37] of intNMF topics top features (ATAC=1000, RNA=300). The colour indicates *−log*_10_ (*p*) for a test of spatial enrichment for a set of regions and a set of genes. The rows are region sets (from ATAC) and the columns are genesets (from RNA). For each region set we test for enrichment at two resolutions: 3000 bp and 100,000 bp to try and capture proximal and distal regulation. (C), (D), (E) show the number of significantly enriched (q *<* 0.05) biological processes’ and disease ontology terms in intNMF topics and MOFA+ factors.

To investigate whether the genes and regions with high weights in a topic are associated with each other, we ran PEGS analysis [37]. This takes sets of genes and genomic regions and tests whether they are mutually enriched, at different spatial resolutions. Generally, the top gene expression features and chromatin accessibility features from a topic are more spatially proximal than expected by chance, as indicated by strong purple diagonal (Figure 3B). This shows strong spatial genomic concordance among corresponding topics in two modalities. Additionally, where there is signal off this diagonal this is often due to topics jointly contributing to the same cell type. For example, topics 10 and 24 both contribute to defining OPCs and topics 11 and 16 both contribute to defining the astrocytes cluster (Figure 2B).

Next we investigated whether the top RNA features of intNMF’s topics defined genes that contribute to particular biological or disease associated processes. We asked whether topic-defined genesets are enriched among identified genesets associated with either biological processes (BP) within the gene ontology (GO) framework or with disease ontologies (DO). We compare the number of enriched GO and DO terms in intNMF topics and MOFA+ topics (Figures 3C, 3D and 3E). intNMF’s genesets have more associated biological processes and disease ontologies than MOFA+ topics, and this advantage is retained irrespective of whether top negative weights, top positive weights or top absolute weights are used in MOFA+ analysis. This indicates that intNMFs topics identify biologically important sets of genes.

### intNMF topics elucidate disease relevant biology

The original study [8] was not able to clearly separate cell types in normal vs Alzheimer’s samples. We therefore asked whether the topics discovered by intNMF could further resolve transcriptional regulatory information about Alzheimer’s disease. Firstly we identified which topics are more strongly activated in Alzheimer’s vs cells from normal samples within the cell types which they are associated with. Figure 4A shows the difference in median topic score (elements of **W**) between cells of the same type from Alzheimer’s and disease free donors. Of the three oligodendrocyte associated topics 19 is more strongly associated with Alzheimer’s whereas 14 and 20 are more strongly associated with normal samples. Additionally, among the two topics associated with astrocytes and two topics associated with microglia there is one topic associated Alzheimer’s (astrocytes topic 11 and microglia topic 18, Figure 4A) and another with control cells (astrocytes topic 16 and microglia topic 8). This can be seen in more detail in Supplementary Figure 9.

**Fig. 4.**
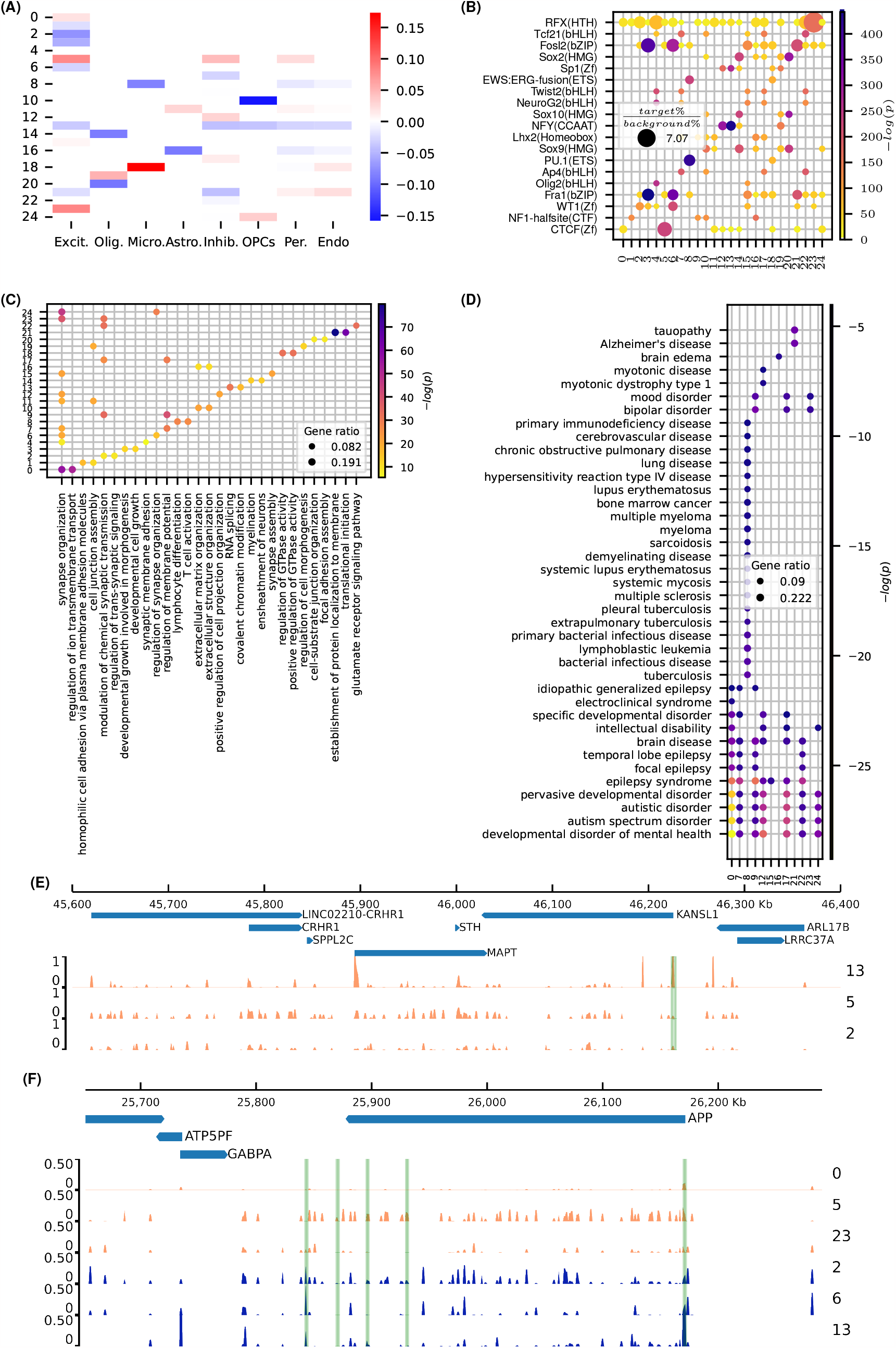
(A) A heatmap showing median topic weight in diseased cell type minus the median topic weight in healthy cell types, value set to zero where topic is not associated with the cell type. (C) Dotplot of biological process GO terms enriched in top RNA features of intNMF’s topics. The top 2 most enriched (by q-value) GO terms are shown. Gene ratio indicates the fraction of top topic genes found in a given geneset. (B) Transcription Factor motif enrichment in top ATAC features of intNMF’s topics. The most enriched (by q value) motif per factor shown. (D) DO ontology enrichment of top RNA features from intNMF topics. Terms are shown if *−log*(*q*) *>* 4. (E) and (F) show genome browser tracks with ATAC feature weights of intNMF topics. (E) shows the *MAPT* locus with a region highlighted which where chromatin accessibility is lost between Alzheimer’s and control samples. (F) shows the *APP* locus with regions enriched for regulatory function (measured by luciferase assay [8]) highlighted in green. All topics shown are associated with excitatory neurons. Orange coloured tracks are Alzheimer’s associated topics and blue coloured tracks are non-Alzheimer’s associated topics.

The top genes in the intNMF topics are associated with GO terms relevant to the cell types associated with the given topic and the Alzheimer’s disease state (Figure 4C). For example the top GO terms for excitatory and inhibitory neuron associated topics 0 and 6, both include “synapse organisation”, and individually, “ion transmembrane transport” is associated with topic 0 whereas topic 6 is associated with “regulation of synapse organization”. Topic 8, associated with non-Alzheimer’s microglia cells, has GO terms associated with leukocyte and T cell activation, in line with their function as the DLPFC’s resident immune cell population, whereas top genes in topic 18 (associated with Alzheimer’s microglia cells), are enriched for genes involved with “positive regulation of GTPase activity”. In addition the top genes of intNMF topics are associated with relevant disease ontologies (DO) (Figure 4D). For example topic 8, associated with microglial cells, shows enrichment of a range of immune system-related diseases. More relevant to potential neuronal cell disfunction, topic 21 is enriched for Alzheimer’s and tauopathy and there are brain-related diseases associated with topics for all cell types.

We also asked whether the chromatin accessibility data from the ATAC-seq features associated with topics gave informative regulatory information. The top ATAC features for topics are enriched for transcription factor (TF) motifs relevant to the cell type the topic is active in. For example, PU.1 is a factor known to be highly expressed in microglia and reduced PU.1 expression is implicated in Alzheimer’s disease onset[38]. It is the significantly enriched in both microglia associated topics (8 and 18), but significantly more so in the non-Alzheimer’s microglia topic (topic 8) (Figure 4B). Other motifs define known master transcription factors of neural-related lineages such as the motif of a key marker of motor neuron and oligodendrocyte differention, olig2, which is enriched in the top accessible regions of two excitatory neuron defining topics (4 and 15).

Finally, we investigated genomic regions analysed in the original paper by looking at the chromatin accessibility topic scores of intNMF topics. The *MAPT* gene codes for the protein tau, and is central to the neuropathology of Alzheimer’s disease [39]. In the original paper they identify its promoter as a region which loses accessibility. This is captured by intNMF’s topics as the region is accessible in topic 13 (an non-Alzheimer’s excitatory cell topic) but is inaccessible in topic 5 (an Alzheimer’s excitatory cell topic, Figure 4E). *APP* encodes amyloid precursor protein, another protein central to Alzheimers pathogenesis. The *APP* locus can be seen in Figure 4F. The highlighted regions are regions found to have regulatory potential via a luciferase assay. The left-most highlighted region exhibited silencer-like properties and, is accessible in the non-Alzheimer’s defining topics in excitatory neurons (blue tracks) but not the ones associated with Alzheimer’s (orange tracks). This suggests a lost repressive regulatory element during Alzheimer’s pathogenesis, although this is paired with a gain in accessibility of *APP*’s promoter suggesting changes in regulation are more complicated than a single lost regulatory element.

These results demonstrate that intNMF topics can identify cell types and subtle differences between disease and normal cells of the same type. Furthermore intNMF’s topic definitions are biologically interpretable and identify important molecular features associated with cell and sub-cell types.

## Discussion

Here we present intNMF, a computationally efficient NMF-based tool for integration of singe cell paired gene expression and chromatin accessibility data. The presented method is benchmarked against other popular methods for single cell data integration. intNMF shows superior clustering performance as measured by ARI and comparable clustering performance as measured by MI to other methods when using the two different sized datasets used in the benchmarking analysis. intNMF and MOFA+ show superior performance to other methods as measured by Silhouette Score, the metric quantifying the structure. However, intNMF has the further advantage over other benchmarked matrix factorisation methods of having significantly faster run time. This is excepting MOJITOO which runs quickly and is a matrix factorisation method but learns canonical components (equivalent of topics), which aren’t linked to the original feature space, but instead to latent representations of the original feature space, trading off some interpretability for speed. In the presented benchmarking work, existing methods which require GPUs for reasonable run times were not applied *e*.*g*. MOWGLI and scVI.

Having determined intNMFs suitability via benchmarking we next performed an analysis on an example dataset of 100, 000 cells from DLPFC from Alzheimer’s positive and negative donors [8]. Through further analysis of intNMF output, we demonstrate that the discovered topics resolve the original cell types identified in the study and link them to relevant molecular features. Additionally the flexibility and granularity of topics as opposed to overall clustering helps elucidate subtle differences in transcriptional regulation between Alzheimer’s and non-Alzheimer’s cells. In this analysis we compared intNMF to one of the best performing existing method, MOFA+, and highlight how the difference in the underlying models, PCA based vs NMF based, can lead to topics which are more cell type-specific and identify important biological features.

intNMF is implemented with an l2 loss, this is a practical decision (similar to MOFA+’s recommendation to use a Gaussian loss for large datasets) due to the efficient update schemes available given an l2 loss. Other cost functions such as the Kullback-Leibler divergence or l1/Poisson could also be applied, which are likely to attend less to highly expressed genes ([40, 41]). Additionally there are flavours of NMF with probabilistic interpretations (primarily by adding sum to one constraints for each cell and each feature) [42], which may further enhance the interpretability of the models. Inference schemes using EM (Expectation Maximisation) or VI (Variational Inference) for updates could be implemented, but they are usually impractical for large datasets. For example, a recent scRNA-seq NMF based method implements an EM algorithm but favours the use of ADAM (a gradient descent based method) due long run times [40]. A potential extension for intNMF, given its speed, is iterative learning -*e*.*g*. fit NMF, add additional topics, remove topics or even fitting topics to subsets of the data. There are existing algorithms which can speed up the process of fitting an NMF model with additional or fewer topics given an NMF model already fit to the data [43]. These techniques could also be utilised to speed up intNMFs model selection via BIC.

intNMF shows best performance (by the metrics presented here) when run with l1-regularisation and a number of topics roughly equal to or slightly more than the true number of different cell types. Use of l1-regularisation helps to induce sparsity in **W** and **H** matrices, which better reflects single cell ‘omics data. It also helps ensure that meaningful features are identified in the feature loading matrices (**H**).

All the presented applications here are of paired chromatin accessibility and gene expression data, however with minor modifications to preprocessing intNMF could be applied to other bi-modal types of data *e*.*g*. paired protein and RNA expression data or spatial gene expression with H&E images. Furthermore with a slight change to the updates of **W** intNMF could be applied to tri-modal datasets *i*.*e* chromatin accessibility, gene expression and protein expression. Given the the ability of intNMF to identify important sets of genes and accessible chromatin regions in populations of cells a further avenue of development for intNMF could be using the topics for regulatory network inference as has been done with regNMF [16] or biological process discovery as has been done recently with Spectra (a method developed for scRNA-seq) [40].

In conclusion intNMF is a fast and accurate method for analysis of single cell multiome data which provide rich low-dimensional representation, facilitates custom downstream analysis (we have shown a flavour in this study) and provides avenues for methodological extensions for further combinations of single cell data modalities.

