## Supplementary material for "Scalable joint non-negative matrix factorisation for paired single cell gene expression and chromatin accessibility data"

September 25, 2023

### 1 Competing methods

#### 1.1 MOFA

MOFA was run in R (R version 4.1.0, MOFA version 1.2.2 and Seurat version 4.2.0) [1]. The preprocessed h5ad file was converted into a Seurat object. MOFA was applied broadly as described in the following tutorial. Seurat's 'FindVariableFeatures' command was used to select genes (N = 5000) and the 'FindTopFeatures' function was used to select accessible regions. MOFA can be run on modalities which have already had their dimensions separately reduced ,i.e., PCA on gene expression matrix and LSI on accessibiility matrix. This improves run time, but makes interpretation harder. We ran MOFA on the original features, having selected the highly variable ones.

#### 1.2 Seurat WNN

Seurat was run in R (R version 4.1.0 and Seurat version 4.2.0) [2]. The preprocessed h5ad file was converted into a Seurat object. Seurat WNN was applied broadly as described in the following tutorial

For preprocessing, instead of normalising then log transforming the data, the SCTransform function was used. Seurat's 'FindVariableFeatures' command was used to select genes (N = 5000) and the 'FindTopFeatures' function was used to select accessible regions. Dimensionality reduction was carried out on the RNA and ATAC matrices via PCA and SVD, respectively. These low dimensional embeddings were then used for WNN using the 'FindMultiModalNeighbors' function. The first component of SVD was removed as it is usually associated with read depth. Leiden clustering and generation of UMAP was then carried out using Seurats tools. This was done due to the difficulty in transferring Seurat's graph representation from Seurat to scanpy/muon.

#### 1.3 scAI

scAI [3] is an NMF based method, which aggregates the epigenomic signals from multiple cells to overcome sparsity. scAI was run in R (R version 4.1.0, seurat version 4.2.0, scAI version 1.0). It was run broadly inline with the following tutorial. However to speed up the process feature selection was carried out using the same method as Seurat WNN and MOFA+.

#### 1.4 scReg

scReg [4] is another NMF based method. It factorises three matrices: gene expression, chromatin accessibility and a *cis*-regulatory potential (accessibility*expression/distance ). scReg was run in R (R version 4.1.0, seurat version 4.2.0, scReg version 0.1.0). scReg was carried out as outlined in it's github readme.

#### 1.5 MOJITOO

MOJITOO implements canonical correlation analysis on modalities which have already had their dimensions reduced [5]. MOJITOO was run in R (R version 4.1.0, seurat version 4.2.0, MOJITOO version 1.0). MOJITOO was carried out as outlined in the tutorial. Seurat's 'FindVariableFeatures' command was used to select genes (N = 3000) and the 'FindTopFeatures' function was used to select accessible regions (min.cutoff = 'q0'). Dimensionality reduction was carried out on the RNA and ATAC matrices via PCA and SVD, respectively. The top 50 components of these low dimensional embeddings were then used for MOJITOO using the 'mojitoo' function. The first component of SVD was removed as it is usually associated with read depth.

### 2 NMF Background

Nonnegative matrix factorisation (NMF) is a linear dimensionality reduction method, similar to PCA. PCA can be considered to minimise the reconstruction error of a matrix while constraining the lower dimensional matrices to contain orthogonal basis. Whereas NMF approximates a nonnegative matrix as the product of two lower dimensional (or low rank) nonnegative matrices which minimise the reconstruction error, *i.e.*

$$\mathbf{X} \approx \mathbf{W}\mathbf{H}. \tag{1}$$

In the case of a scRNA-seq dataset: $\mathbf{X} \in \mathbb{R}_+^{M \times N}$ is a nonnegative sample by gene matrix to be factorised ($M$ is the number of cells and $N$ is the number of genes), the columns of $\mathbf{W} \in \mathbb{R}_+^{M \times K}$ contain the nonnegative coordinates of each cell in terms of the nonnegative basis vectors given by the rows of $\mathbf{H} \in \mathbb{R}_+^{K \times N}$. $K$ is the desired number of dimensions for the factorisation, referred to here as topics due to NMFs similarity with topic modelling (expand). $K$ is selected such that $K << M$ or $N$. The nonnegativity constraint means the low rank matrices reflect the original matrix in a more interpretable fashion as the basis vectors learn a parts based representation of the original data [6] and they contain real positive numbers like the original data. For example, when NMF is applied to a collection of images each basis vector is an image.

$\mathbf{W}$ and $\mathbf{H}$ can be found by minimising a cost function describing the reconstruction error *e.g.*

$$min_{\mathbf{W},\mathbf{H}} f(\mathbf{W},\mathbf{H}) = ||\mathbf{X} - \mathbf{WH}||_F^2 \quad s.t. \quad \mathbf{W} \geq 0, \mathbf{H} \geq 0 \tag{2}$$

in this case the cost function used is the Frobenius norm,

The multiplicative update method proposed by Li *et al* is popular due to its simplicity to implement and guaranteed nonnegative values at each iteration [7]. However the hierarchical alternating least squares (HALS) and alternating nonnegative least squares (ANLS) methods converge faster in practice [8]. Both of these methods can be explained by the common framework of block coordinate descent (BCD) [9]. ANLS fixes $\mathbf{W}$ or $\mathbf{H}$ sequentially and updates the other. This reduces to a convex nonnegative least squares problem which can be solved efficiently, *i.e.*

$$\begin{aligned} \mathbf{W} \leftarrow argmin_{\mathbf{W} \leq 0} f(\mathbf{W},\mathbf{H}) \text{ and} \\ \mathbf{H} \leftarrow argmin_{\mathbf{W} \leq 0} f(\mathbf{W},\mathbf{H}) \end{aligned} \tag{3}$$

intNMF implements an accelerated version of HALs proposed by Gillis and Gilneur [10]. HALs extends ANLS by updating a single column of $\mathbf{H}$ or $\mathbf{W}$ and fixing all other columns *e.g.*

$$\mathbf{w_k} \leftarrow argmin_{\mathbf{w_k} \geq 0} f(\mathbf{W},\mathbf{H}) \tag{4}$$

where $\mathbf{w_k}$ is the $k^{th}$ column of $\mathbf{W}$. Using the Frobenius norm as the cost function gives:

$$min_{\mathbf{w_k}} ||(\mathbf{X} - \sum_{l \neq k} \mathbf{w_l h_l}) - \mathbf{w_k h_k}||_F^2 = ||\mathbf{R} - \mathbf{w_k h_k}||_F^2 \tag{5}$$

where $\mathbf{h_k}$ is a row of $\mathbf{H}$ and $\mathbf{R}$ is the residual from reconstruction using the other columns of $\mathbf{W}$ and rows $\mathbf{H}$ *i.e.*

$$\mathbf{R} = \mathbf{X} - \sum_{l \neq k} \mathbf{w_l h_l} \tag{6}$$

This gives update equation for $\mathbf{W}$ as

**Algorithm 1** W HALS

**procedure** W_HALS($\mathbf{W}, rnaHHt, rnaMHt, atacHHt, atacMHt, precompute\_time, \alpha = 0.5, \delta = 0.1$)

$i \leftarrow 1$

$start\_time \leftarrow get\_time()$

**repeat**

$\epsilon \leftarrow 0$

**for** k, $\mathbf{w_k} \in enumerate(columns(\mathbf{W}))$ **do**

$\Delta\mathbf{w} \leftarrow max(\frac{rnaMHt[:,k] - \mathbf{W}rnaHHt[:,k] + atacMHt[:,k] - \mathbf{W}atacHHt[:,k]}{rnaHHt[k,k] + atacHHt[k,k]}, -\mathbf{w_k})$

$\mathbf{w_k} \leftarrow \mathbf{w_k} + \Delta\mathbf{w}$

$\epsilon \leftarrow \epsilon + \Delta\mathbf{w}\Delta\mathbf{w}^T$

$\mathbf{w_k}[\mathbf{w_k} == 0] \leftarrow max(\mathbf{w_k}) \times 1e - 16$

**if** $i == 1$ **then**

$\epsilon_0 \leftarrow \epsilon$

$i \leftarrow 0$

$first\_iteration\_time \leftarrow get\_time() - start\_time$

$time\_in\_function \leftarrow get\_time() - start\_time$

**until** $((time\_in\_function > \alpha(first\_iteration\_time + precompute\_time)$ OR $(\epsilon \leq \delta^2\epsilon_0))$

return($\mathbf{W}$)

**Algorithm 2** H HALS

**procedure** H_HALS($\mathbf{H}, WtM, WtW, precompute\_time, \alpha = 0.5, \delta = 0.1$)
    $i \leftarrow 1$
    $start\_time \leftarrow get\_time()$
    **repeat**
        $\epsilon \leftarrow 0$
        **for** k, $\mathbf{h_k} \in enumerate(rows(\mathbf{W}))$ **do**
            $\Delta\mathbf{h} \leftarrow max(\frac{WtM[k,:]-WtW[k,:]\mathbf{H}}{WtW[k,k]}, -\mathbf{h_k})$
            $\mathbf{h_k} \leftarrow \mathbf{h_k} + \Delta\mathbf{h}$
            $\epsilon \leftarrow \epsilon + \Delta\mathbf{h}\Delta\mathbf{h}^T$
            $\mathbf{h_k}[\mathbf{h_k} == 0] \leftarrow max(\mathbf{h_k}) \times 1e - 16$
        **if** $i == 1$ **then**
            $\epsilon_0 \leftarrow \epsilon$
            $i \leftarrow 0$
            $first\_iteration\_time \leftarrow get\_time() - start\_time$
        $time\_in\_function \leftarrow get\_time() - start\_time$
    **until** (($time\_in\_function > \alpha(first\_iteration\_time + precompute\_time)$ OR ($\epsilon \leq \delta^2\epsilon_0$))
    return($\mathbf{H}$)

$$\mathbf{w_k} \leftarrow [\mathbf{w_k} + \frac{(\mathbf{XH^T})_k - (\mathbf{WHH^T})_k}{(\mathbf{HH^T})_{kk}}]_+ \tag{7}$$

where $[.]_+$ indicates $max([.], 0)$. Similarly, the update equation for $\mathbf{H}$ is given by

$$\mathbf{h_k} \leftarrow [\mathbf{h_k} + \frac{(\mathbf{X^TW})_k - (\mathbf{W^TWH})_k}{(\mathbf{W^TW})_{kk}}]_+ \tag{8}$$

When updating $\mathbf{W}$, $\mathbf{XH^T}$ and $\mathbf{HH^T}$ are unaffected by updating columns of $\mathbf{W}$, therefore it is computationally efficient to precompute these values update all columns of $\mathbf{W}$. intNMF applies an accelerated version of the HALs proposed by gillis *et al* where $\mathbf{W}$ and $\mathbf{H}$ are updated multiple times at each epoch. After updating all columns of $\mathbf{W}$ or $\mathbf{H}$ the matrix is updated again if both of the following criteria are met:

1. epoch run time < (precompute time (*e.g.* $\mathbf{X^TW}$ and $\mathbf{H^TH}$) + last iteration duration)/2

2. sum of squares of update to $\mathbf{W}$ or $\mathbf{H}$ at last iteration > 0.01*sum of squares of update to $\mathbf{W}$ or $\mathbf{H}$ at first iteration

Updates to a single matrix are stopped when either of the conditions are no longer met.

### 3 intNMF

intNMF aims to factorise two matrices $(\mathbf{X_1}, \mathbf{X_2})$ so that they share a single $\mathbf{W}$ but have distinct $\mathbf{H}$ matrices, *e.g.*

$$min_{\mathbf{W},\mathbf{H_1},\mathbf{H_2}} ||\mathbf{X_1} - \mathbf{WH_1}||_F^2 + ||\mathbf{X_2} - \mathbf{WH_2}||_F^2 \tag{9}$$

This is achieved by applying the HALs method outlined above. However $\mathbf{W}$ updates must be calculated w.r.t both $\mathbf{X_1}$ and $\mathbf{X_2}$ simultaneously. This results in the same updates for $\mathbf{H_1}$ and $\mathbf{H_2}$. However the update for $\mathbf{W}$ is slightly different. We have the minimisation problem:

$$min_{\mathbf{w_k}} ||\mathbf{h_k^{RNA}} \mathbf{w_k^T} - (\mathbf{R_k^{RNA}})^\mathbf{T}||_F^2 + ||\mathbf{h_k^{ATAC}} \mathbf{w_k^T} - (\mathbf{R_k^{RNA}})^\mathbf{T}||_F^2 \tag{10}$$

where $\mathbf{R_k^{RNA}} \in \mathbb{R}_+^{M\times N}$, $\mathbf{h_k^{RNA}} \in \mathbb{R}_+^N$, $\mathbf{R_k^{ATAC}} \in \mathbb{R}_+^{M\times L}$, and $\mathbf{h_k^{ATAC}} \in \mathbb{R}_+^L$ are given. This can be further broken down by taking the sum for each sample *i.e.*

$$\sum_{n=1}^{N} min_{w_{k,n}} ||\mathbf{h_k}^{RNA} w_{k,n} - \mathbf{r^{RNA}_{k,n}}||_2^2 + ||\mathbf{h^{ATAC}_k} w_{k,n} - \mathbf{r^{RNA}_{k,n}}||_F^2 \tag{11}$$

Expanding this we get:

$$\begin{aligned} h(w_{k,n}) = ||\mathbf{h^{RNA}_k}||_2^2 w_{k,n} - 2w_n \mathbf{h_k} \mathbf{r^{RNA}_{k,n}} + ||\mathbf{r^{RNA}_{k,n}}||_2^2 + \\ ||\mathbf{h^{ATAC}_k}||_2^2 w_{k,n} - 2w_n \mathbf{h_k} \mathbf{r^{ATAC}_{k,n}} + ||\mathbf{r^{ATAC}_{k,n}}||_2^2 \end{aligned} \tag{12}$$

taking the derivative with respect to $w_{k,n}$ gives

$$\begin{aligned} \frac{\partial h}{\partial w_{k,n}} = 2w_{k,n} ||\mathbf{h^{RNA}_k}||_2^2 - 2(\mathbf{r^{RNA}_{k,n}})^{\mathbf{T}} \mathbf{h^{RNA}_k} \\ + 2w_{k,n} ||\mathbf{h^{ATAC}_k}||_2^2 - 2(\mathbf{r^{ATAC}_{k,n}})^{\mathbf{T}} \mathbf{h^{ATAC}_k} \end{aligned} \tag{13}$$

This gives an update for $w_{k,n}$ as

$$w_{k,n} = \frac{[(\mathbf{r^{RNA}_{k,n}})^{\mathbf{T}} \mathbf{h^{RNA}_k}]_+ + [(\mathbf{r^{ATAC}_{k,n}})^{\mathbf{T}} \mathbf{h^{ATAC}_k}]_+}{||\mathbf{h^{RNA}_k}||_2^2 + ||\mathbf{h^{ATAC}_k}||_\mathbf{2}^\mathbf{2}} \tag{14}$$

And in turn the update for $\mathbf{w_k}$ is given by:

$$\mathbf{w_k} = \frac{[\mathbf{R^{RNA}_k} \mathbf{h^{RNA}_k}]_+ + [(\mathbf{R^{ATAC}_k})^{\mathbf{T}} \mathbf{h^{ATAC}_k}]_+}{||\mathbf{h^{RNA}_k}||_2^2 + ||\mathbf{h^{ATAC}_k}||_\mathbf{2}^\mathbf{2}} \tag{15}$$

In practice it is useful to replace $\mathbf{R_k}$ so that it does not need to be calculated each iteration.

$$\mathbf{w_k} = \frac{[(\mathbf{X_{RNA}} + \mathbf{h^{RNA}_k} \mathbf{w_k} - \mathbf{WH}) \mathbf{h^{RNA}_k} + (\mathbf{X_{ATAC}} + \mathbf{h^{ATAC}_k} \mathbf{w_k} - \mathbf{WH_{ATAC}}) \mathbf{h^{ATAC}_k}]_+}{||\mathbf{h^{RNA}_k}||_2^2 + ||\mathbf{h^{ATAC}_k}||_\mathbf{2}^\mathbf{2}} \tag{16}$$

$$\mathbf{w_k} = \mathbf{w_k} + \frac{[\mathbf{X_{RNA}} \mathbf{h^{RNA}_k} - \mathbf{WH_{RNA}} \mathbf{h^{RNA}_k} + \mathbf{X_{ATAC}} \mathbf{h^{ATAC}_k} - \mathbf{WH_{ATAC}} \mathbf{h^{ATAC}_k}]_+}{||\mathbf{h^{RNA}_k}||_2^2 + ||\mathbf{h^{ATAC}_k}||_\mathbf{2}^\mathbf{2}} \tag{17}$$

Or written in a more similar style to the updates as they appear in intNMFs python implementation:

$$\mathbf{W}[:, \mathbf{k}] = \mathbf{W}[:, \mathbf{k}] + max(\frac{\mathbf{X_{RNA}}\mathbf{H_{RNA}^{T}}[:, \mathbf{k}] - \mathbf{W}(\mathbf{H_{RNA}}\mathbf{H_{RNA}^{T}}[:, \mathbf{k}])}{(\mathbf{H_{RNA}}\mathbf{H_{RNA}}^{\mathbf{T}}[\mathbf{k}, \mathbf{k}]) + (\mathbf{H_{ATAC}}\mathbf{H_{ATAC}^{T}}[\mathbf{k}, \mathbf{k}])} + \frac{\mathbf{X_{ATAC}}\mathbf{H_{ATAC}^{T}}[:, \mathbf{k}] - \mathbf{W}(\mathbf{H_{ATAC}}\mathbf{H_{ATAC}^{T}}[:, \mathbf{k}])}{(\mathbf{H_{RNA}}\mathbf{H_{RNA}^{T}}[\mathbf{k}, \mathbf{k}]) + (\mathbf{H_{ATAC}}\mathbf{H_{ATAC}^{T}}[\mathbf{k}, \mathbf{k}])}, -\mathbf{W}[:, \mathbf{k}]) \quad (18)$$

The updates for l1-regularisation can be found by substituting an elementwise l1 penalty (for $\mathbf{W}$, $\mathbf{H_{atac}}$ and $\mathbf{H_{rna}}$) into equation 9 and following a similar procedure.

### 4 Results

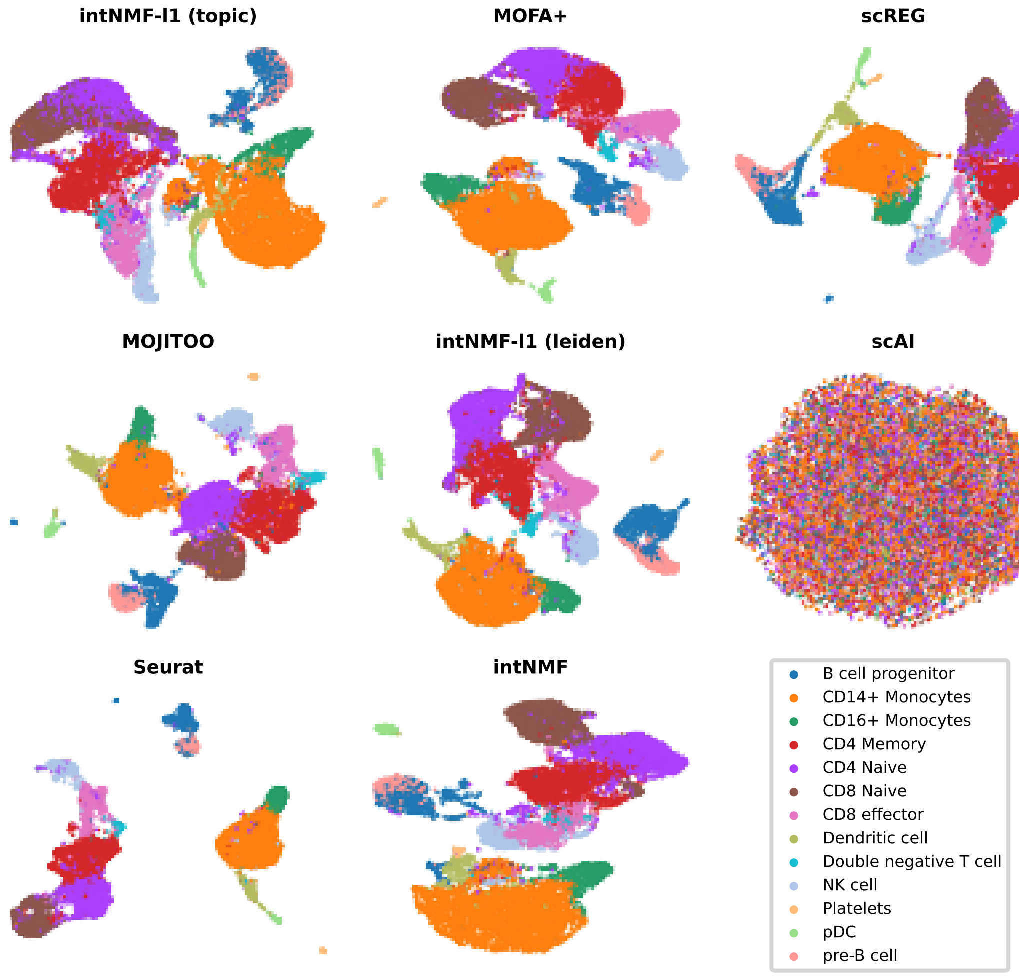


**Figure 1:** UMAPS generated from joint embeddings produced by benchmark methods on the PBMC-Multiome dataset. Cells coloured by cell type annotations.

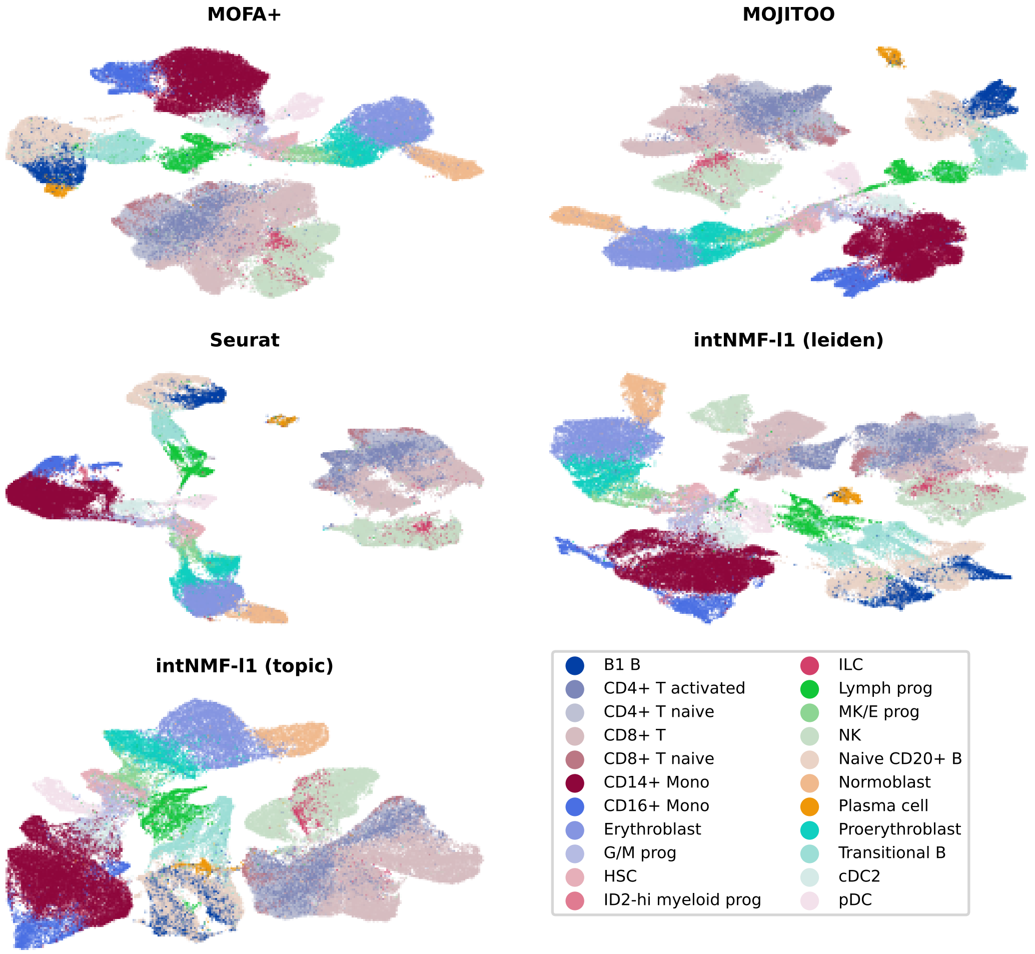


**Figure 2:** UMAPS generated from joint embeddings produced by benchmark methods on the BMMC-Multiome dataset. Cells coloured by cell type annotations.

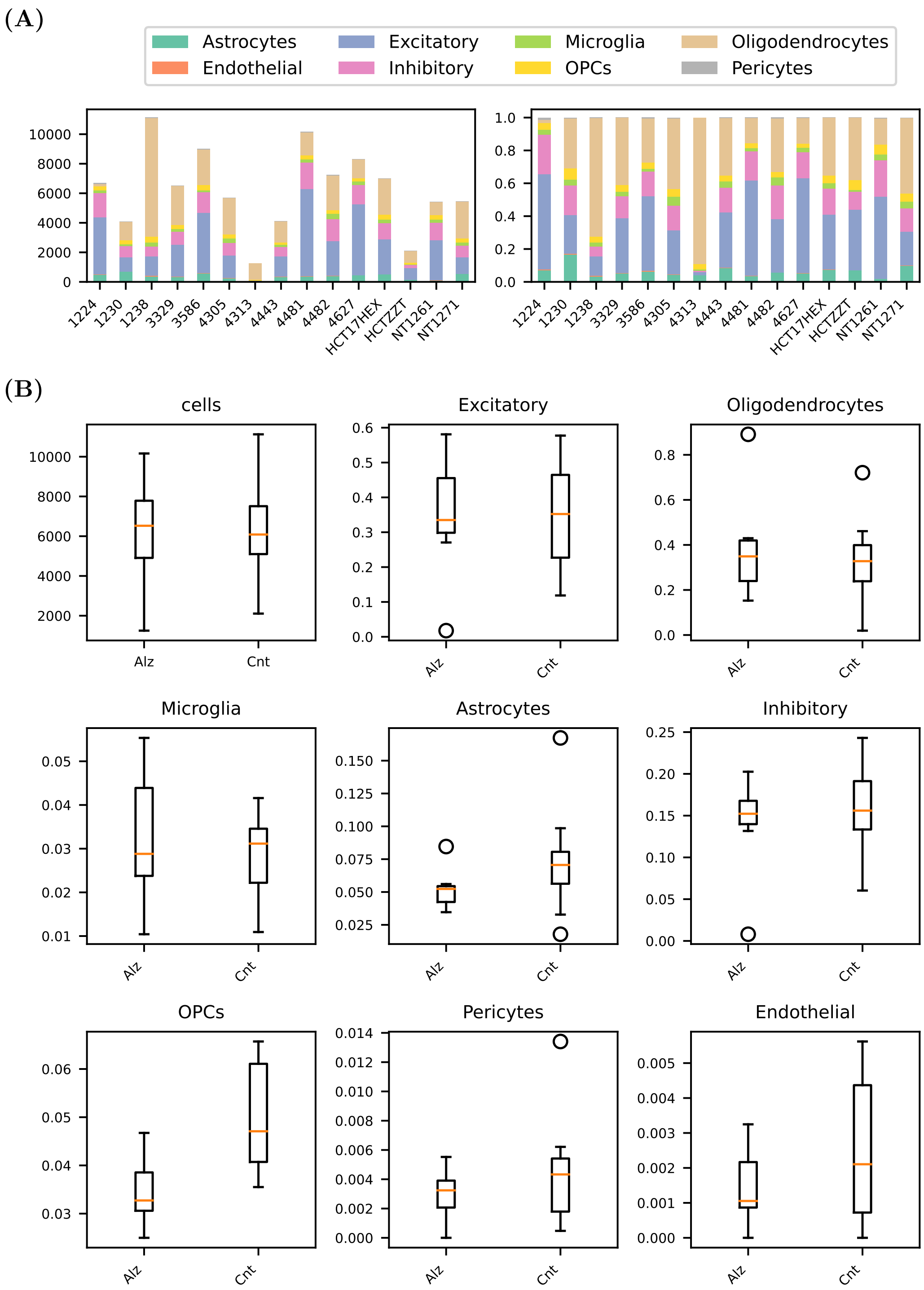


**Figure 3:** Summary plots for Alzheimers-multiome dataset. (A) Cell number (left) and proportion (right) by sample. (B) box plots of proportion of cell types by Alzheimer's status

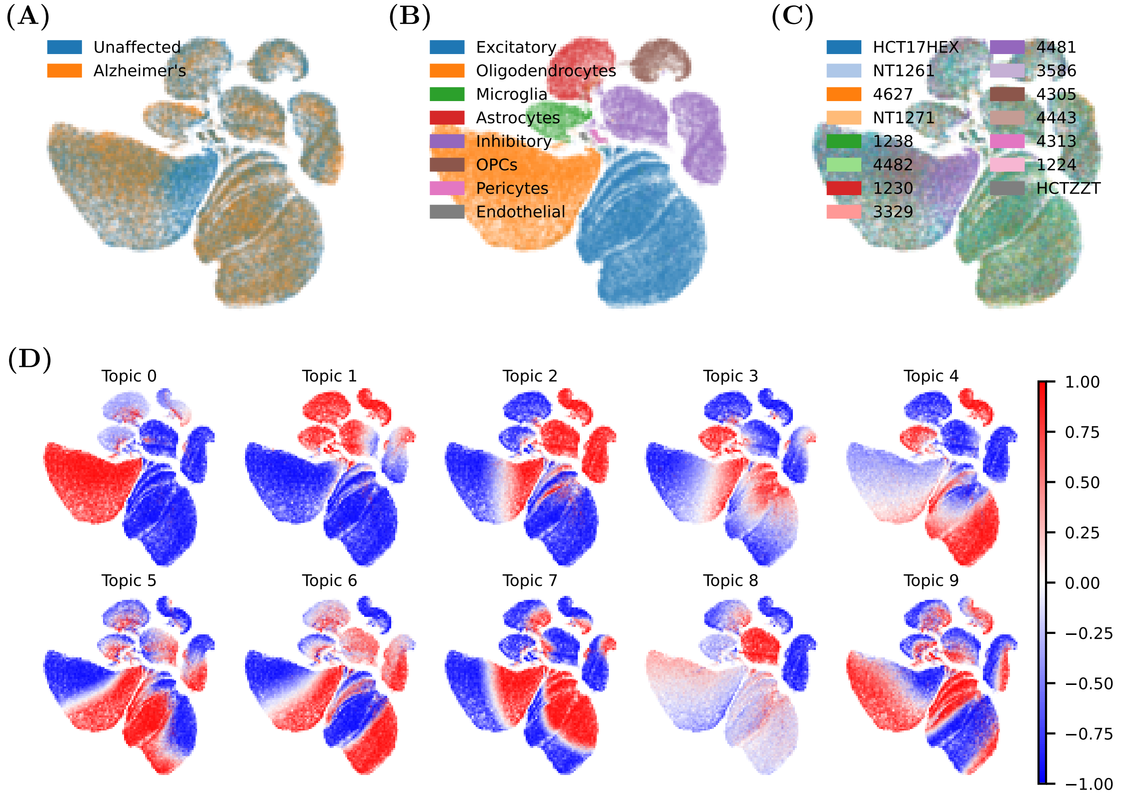


**Figure 4:** Alzheimers dataset UMAPs generated from MOFA+ embedding coloured by disease status (A), cell type (B), sample id (C) and factor weights (D).

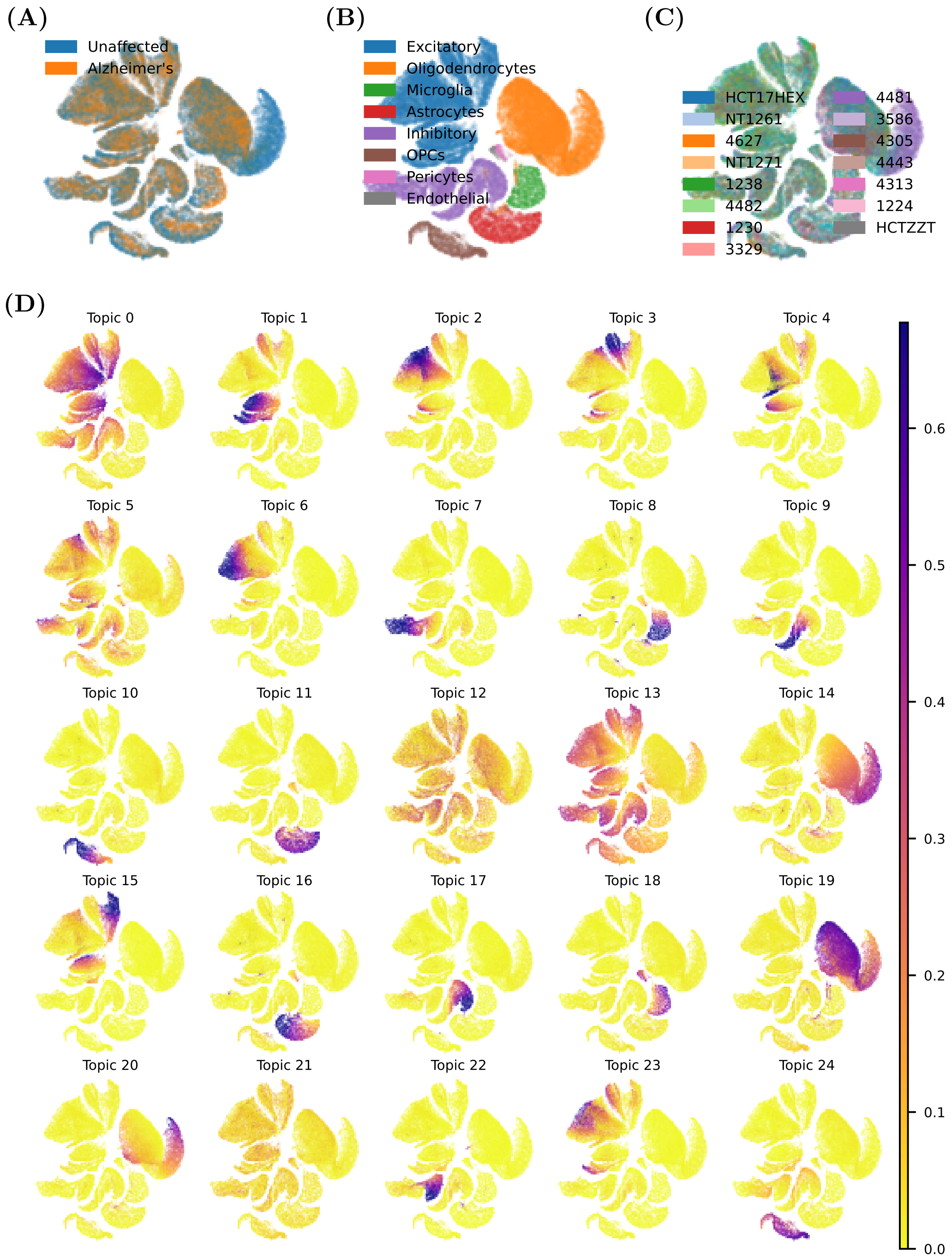


**Figure 5:** Alzheimers dataset UMAPs generated from intNMF embedding coloured by disease status (A), cell type(B), sample id (C) and factor weights (D).

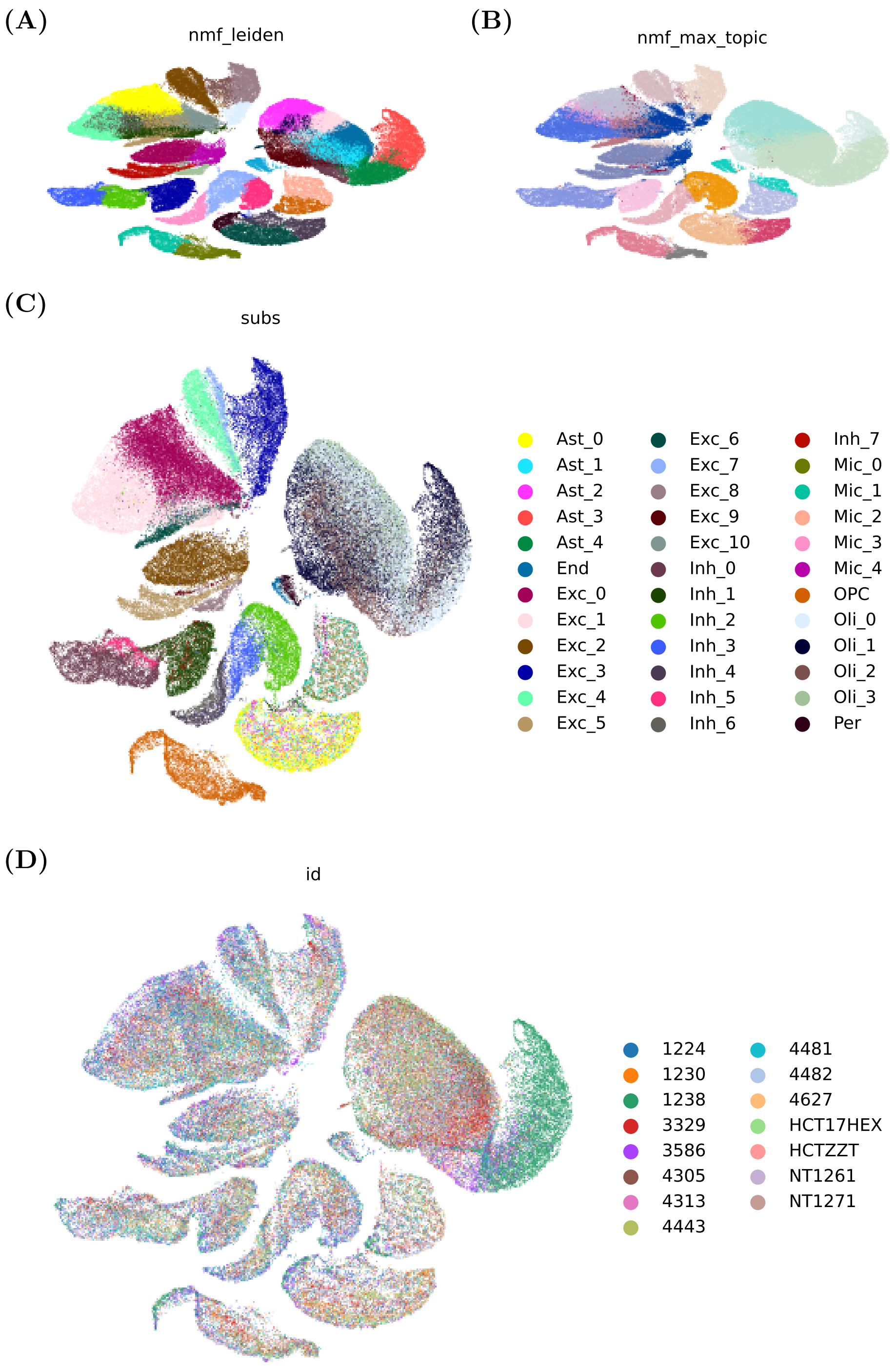


**Figure 6:** Alzheimers dataset UMAPs generated from intNMF embedding (**W**) coloured by Leiden clusters from intNMF embedding (A), intNMF max topic (B), sub cell type annotations (C) and sample id(D).

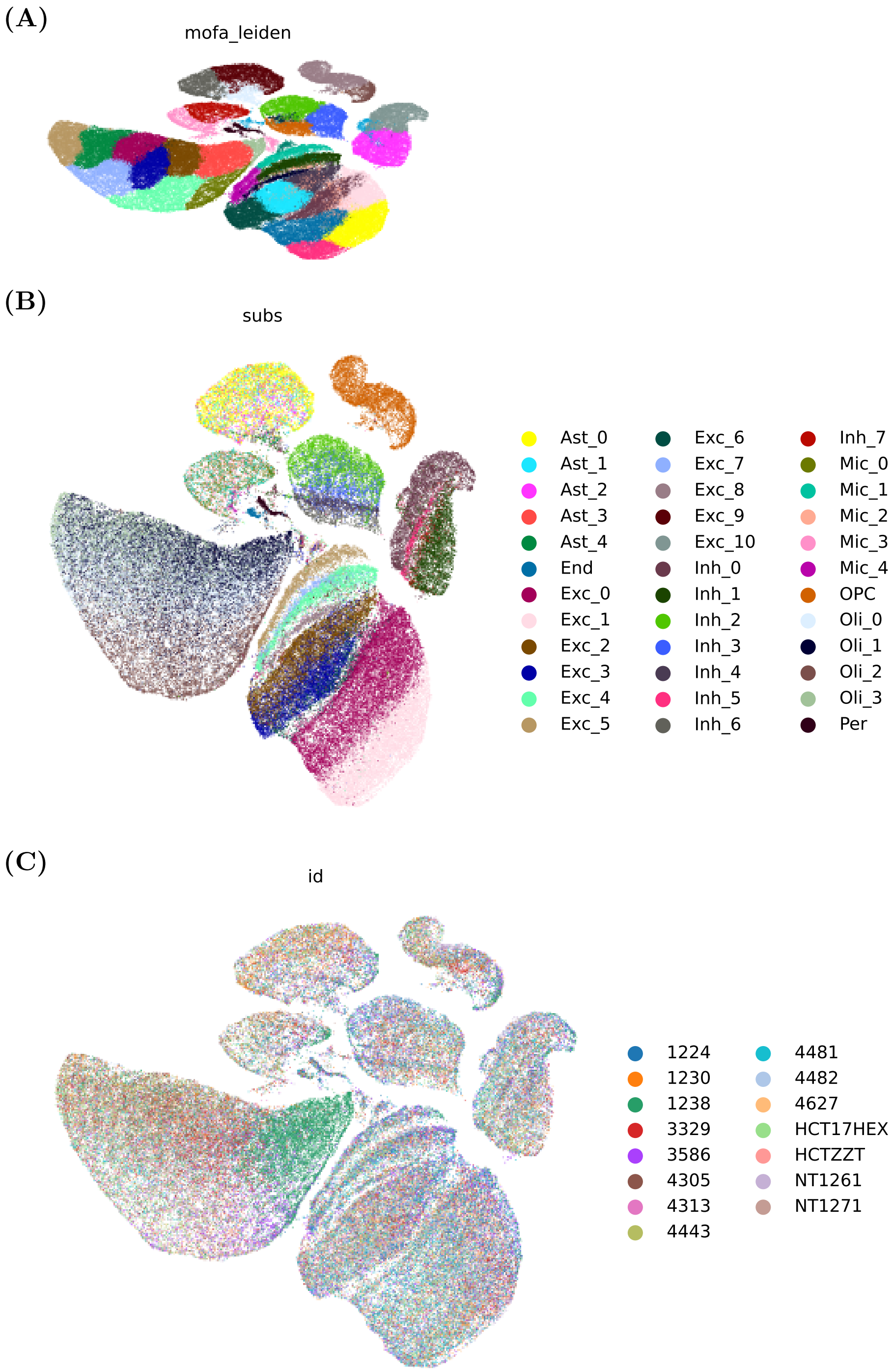


**Figure 7:** Alzheimers dataset UMAPs generated from MOFA embedding coloured by Leiden clusters generated from MOFA embedding (A), sub cell type annotations (B) and sample id (C).

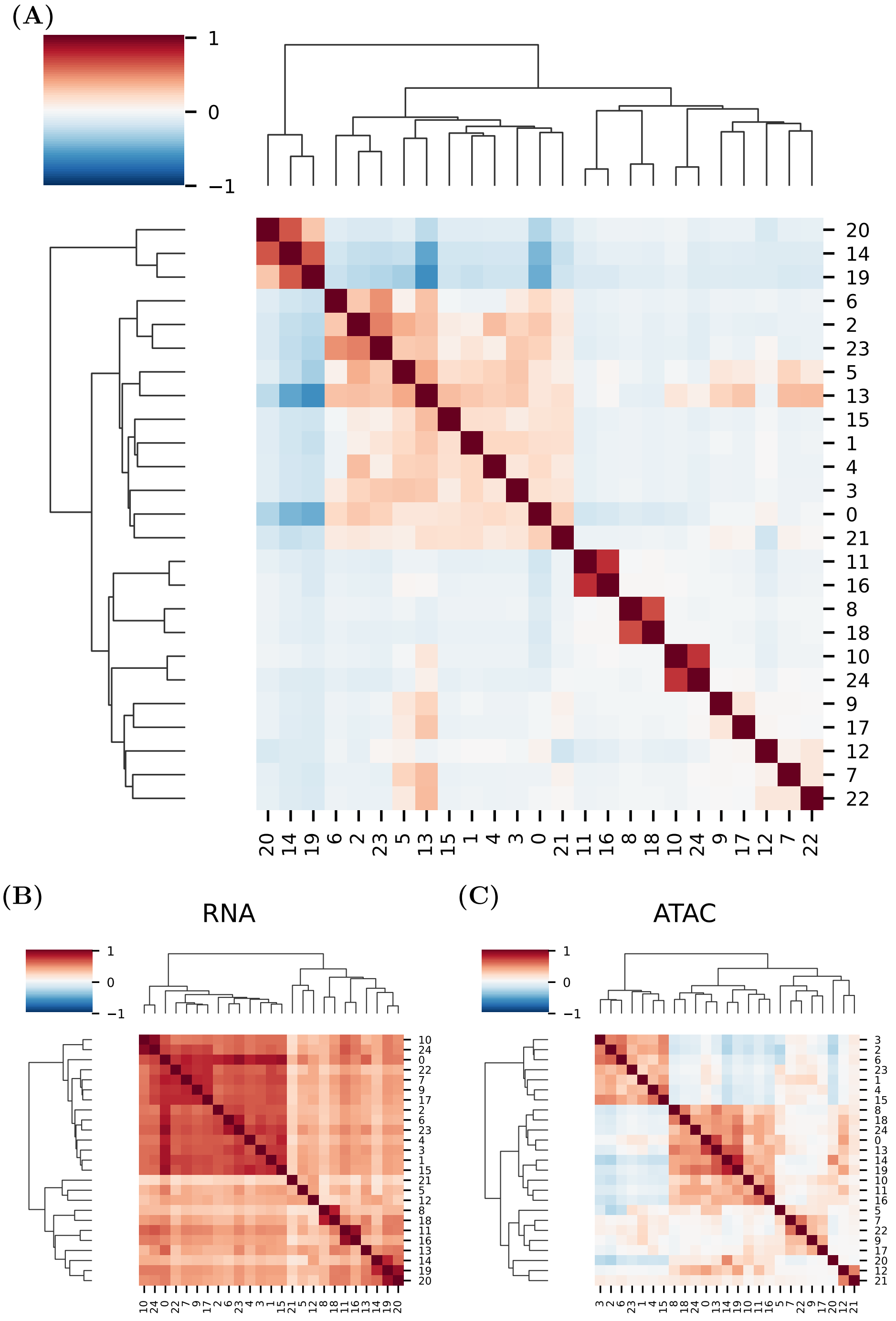


**Figure 8:** Heatmaps showing correlations of topics (all plots are topic x topic). (A) shows correlation based on topic scores *i.e.* $\mathbf{W}$. (B) shows correlations based on RNA loadings *i.e.* $\mathbf{H}_{\mathbf{RNA}}$ and (C) shows correlations based on ATAC loadings *i.e.* $\mathbf{H}_{\mathbf{ATAC}}$

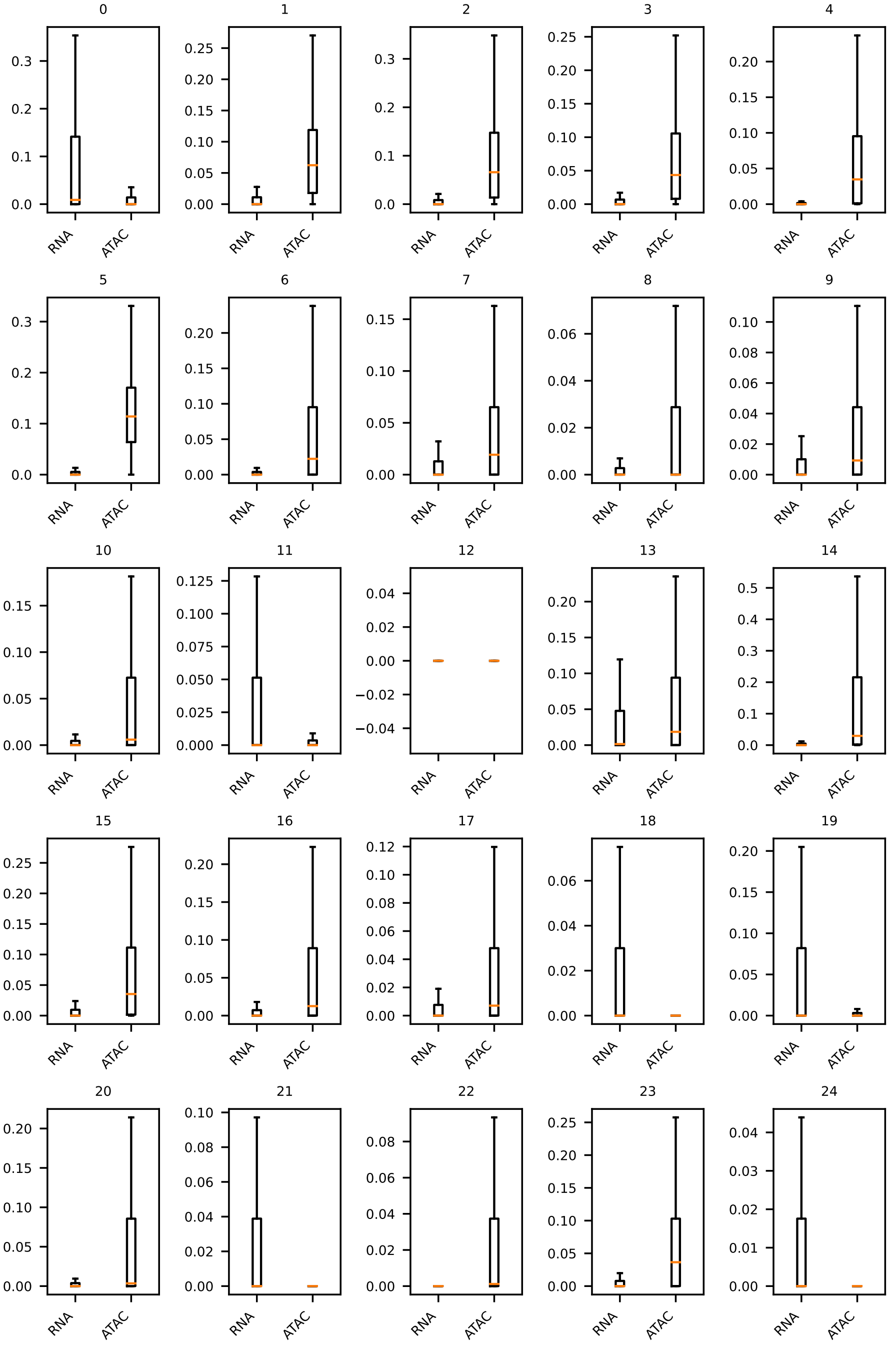


**Figure 9:** Boxplots of ATAC and RNA loading scores (*i.e.* rows of $\mathbf{H_{RNA}}$ and $\mathbf{H_{ATAC}}$) for all topics for intNMF model fit on the Alzheimer-multiome dataset.

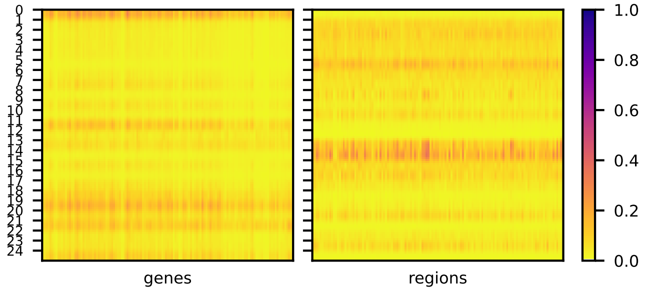


**Figure 10:** Heatmap of RNA and ATAC loading matrices (*i.e.* $\mathbf{H_{RNA}}$ left and $\mathbf{H_{ATAC}}$ right)

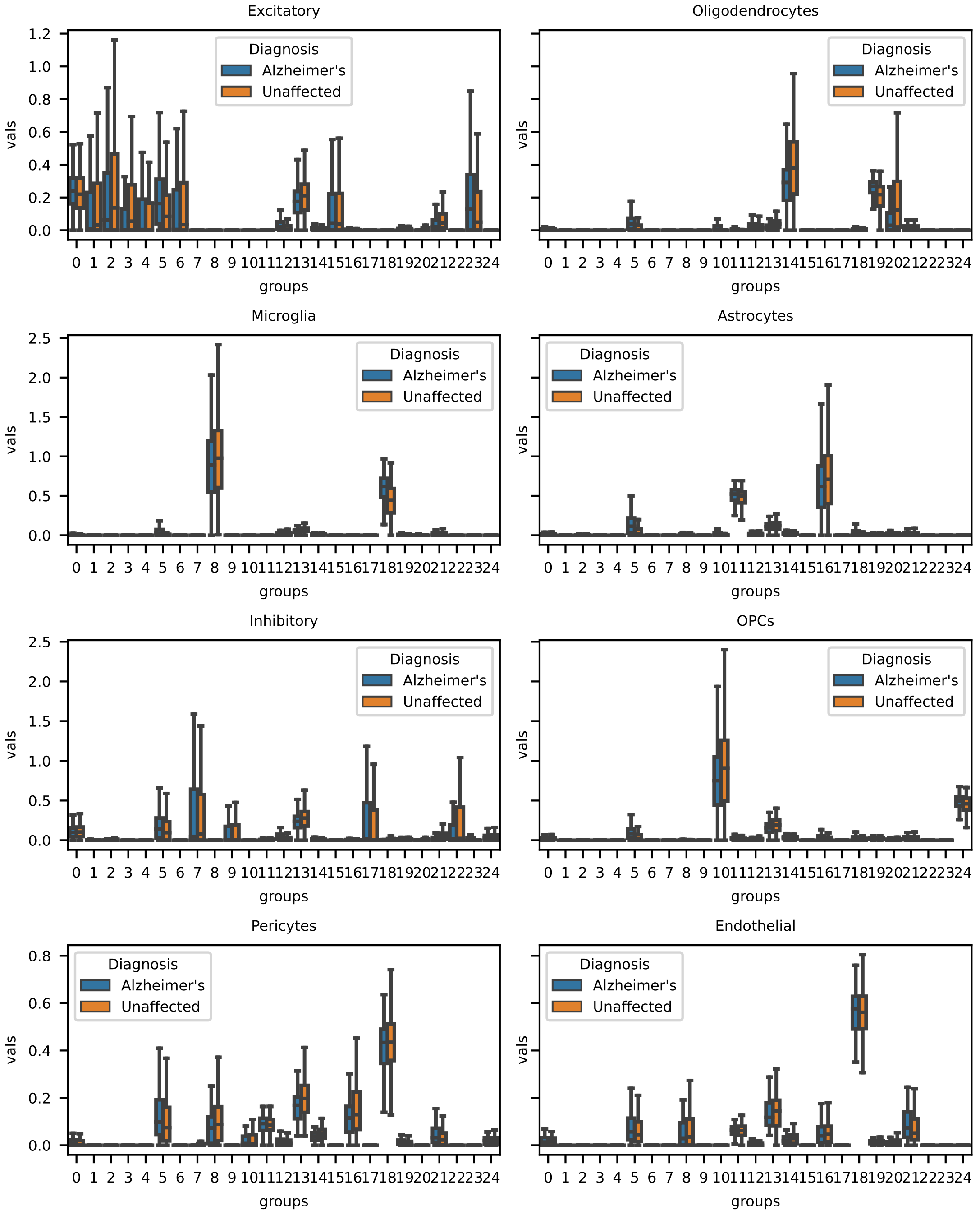


**Figure 11:** Box plots of topic scores ($\mathbf{W}$) in cell sub-populations grouped by disease status: Alzheimer's and Unaffected.

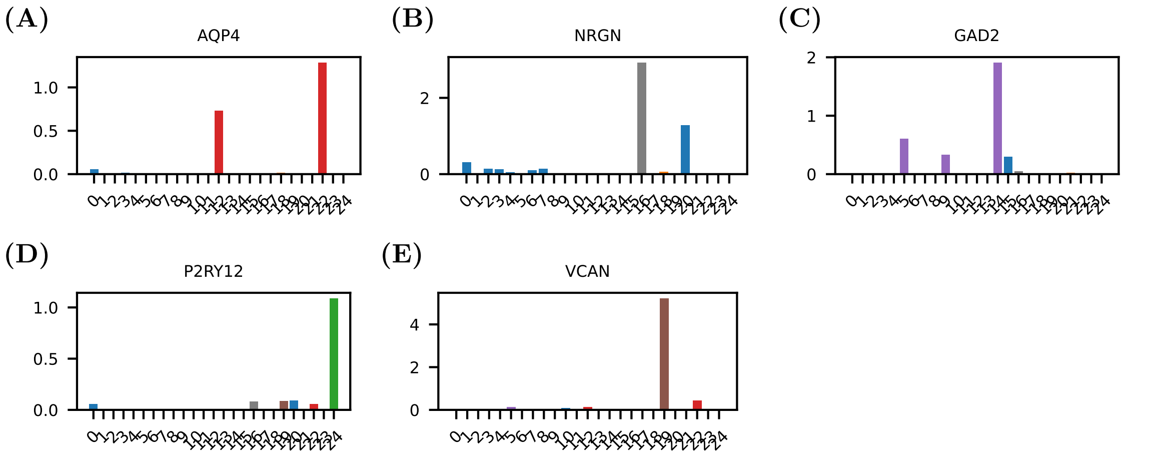


**Figure 12:** Topic weights of cell markers, topics colours by cell type they are associated with

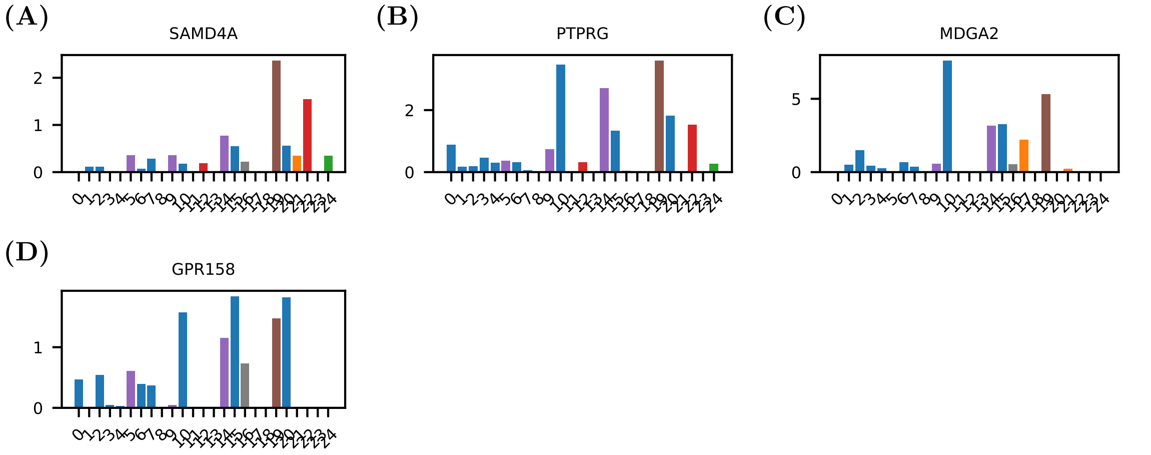


**Figure 13:** Topic weights of genes differentially expressed between Alheimers and non-diseased samples, topics coloured by cell type they are associated with.
